# metaPR^2^: a database of eukaryotic 18S rRNA metabarcodes with an emphasis on protists

**DOI:** 10.1101/2022.02.04.479133

**Authors:** Daniel Vaulot, Clarence Wei Hung Sim, Denise Ong, Bryan Teo, Charlie Biwer, Mahwash Jamy, Adriana Lopes dos Santos

**Affiliations:** UMR 7144, ECOMAP, CNRS, Sorbonne Université, Station Biologique de Roscoff, 29680 Roscoff, France; Asian School of the Environment, Nanyang Technological University, 50 Nanyang Avenue, Singapore 639798; Department of Organismal Biology (Systematic Biology), Uppsala University, Uppsala, Sweden

**Keywords:** 18S rRNA, metabarcodes, database, R, shiny, PCR, protists

## Abstract

In recent years, metabarcoding has become the method of choice for investigating the composition and assembly of microbial eukaryotic communities, and an increasing number of environmental datasets are being published. Although unprocessed sequence files are often publicly available, processed data, i.e. sequences clustered as operational taxonomic units (OTUs) or amplicon sequence variants (ASVs) are rarely at hand in a comparable format. This hampers comparative studies between different environments and datasets, for example examining the biogeographical patterns of specific groups/species, as well analysing the micro-genetic diversity within these groups. Here, we present a newly-assembled database of processed 18S rRNA metabarcodes that are annotated with the PR^2^ reference sequence database. This database, called metaPR^2^, contains 41 datasets corresponding to more than 4,000 samples and 73,000 ASVs. The database is accessible through both a web-based interface (https://shiny.metapr2.org) and as an R package, and should prove very useful to all researchers working on protist diversity in a variety of systems.

## Introduction

Protists, i.e. microbial eukaryotes that are not plants, animals or fungi (Archibald et al. 2017), are one of the most dominant life forms on earth, comprising up to 80% of the total eukaryotic diversity in the environment (De Vargas et al. 2015; Mahé et al. 2017; Massana et al. 2015). Protists play key ecological roles and are involved in primary productivity, nutrient cycling and carbon sequestration. It is thus crucial to assess protist diversity and the factors that determine community composition in order to predict how protists will respond to environmental change (Cavicchioli et al. 2019). While protists have historically been more difficult to study due to their small size, the explosion of metabarcoding studies over the past ten years have greatly expanded our knowledge of these organisms (Burki et al. 2021; Santoferrara et al. 2020).

Metabarcoding reveals the taxa present in an environment by amplifying and then massively sequencing a standardised genetic marker (Santoferrara 2019; Taberlet et al. 2012). In recent years, it has become a very powerful and widespread approach to investigate protist diversity in a range of environments (marine, freshwater, soils, microbiomes etc.). By far, the most common marker used for eukaryotic microbes is the gene coding for small ribosomal subunit RNA (18S rRNA). This gene has the advantage of being universal and having well annotated reference databases such as Silva or PR^2^ (Guillou et al. 2013; Quast et al. 2013) which allow, for many protist groups, a precise taxonomic assignation. Within the 18S rRNA gene, several variable regions have been used as barcodes, in particular the V4 region located near the middle of the gene and the shorter V9 region located at its 3’ end (Burki et al. 2021; Pawlowski et al. 2012). The V4 region in particular is currently most often used in recent studies (Lopes dos Santos et al. 2021). Over the years, metabarcoding has been used to study various aspects of protist diversity. The first studies aimed to simply establish the real extent of eukaryotic diversity that was underestimated with traditional clone library approaches (e.g. Stoeck et al. 2009). In marine waters, metabarcoding studies extended quickly and now tackle more focused questions, for example analysing the distribution of protist groups in the ocean as a function of their size (De Vargas et al. 2015), the diversity of heterotrophic protists in the deep layers of the ocean (Giner et al. 2020; Obiol et al. 2021), detailed biogeographic distribution of specific taxa (e.g. Malviya et al. 2016; Yau et al. 2020), factors structuring marine plankton communities (Logares et al. 2020; Sommeria-Klein et al. 2021), and the seasonal succession of taxa (e.g. Giner et al. 2019; Lambert et al. 2019). Fewer metabarcoding studies have been carried out in freshwater and soils, but that is rapidly changing with some large scale studies (e.g. for soils Mahé et al. 2017).

Most eukaryotic metabarcoding studies have targeted one specific environment, thereby preventing large scale comparisons. On the other hand, large metabarcoding projects using the 16S rRNA gene have been undertaken such as the Earth Microbiome Project which encompassed more than 23,000 samples of both free-living or host-associated microbes, and allowed inferences of global patterns of prokaryotic diversity (Thompson et al. 2017). For eukaryotic 18S rRNA, large expeditions for sample collections have been undertaken in particular for marine systems such as *Tara* Oceans, Ocean Sampling Day (OSD) and Malaspina (De Vargas et al. 2015; Duarte 2015; Kopf et al. 2015). Many studies that performed analyses on the global ocean microbiota have used one or several of these three datasets, in particular *Tara* Oceans (e.g. Ibarbalz et al. 2019; Sommeria-Klein et al. 2021). Many more smaller-scale metabarcoding studies have also been carried out, in particular for environments that have been not sampled by these expeditions, such as soils or freshwater lakes and rivers (Lopes dos Santos et al. 2021). Unfortunately, it is difficult to combine the data from these studies with those of the large scale expeditions for a range of reasons. First, even if the unprocessed data files containing raw reads have been deposited to GenBank SRA (Small Reads Archive), secondary data, i.e. clustered sequences at a certain similarity level, so-called Operational Taxonomic Units (OTUs), or Amplified Sequence Variants (ASVs, Callahan et al. 2016) that do not depend on a specific similarity threshold, are rarely available or, if available, hard to locate since they are stored in a range of formats (DOCX, XLSX or TXT files) as supplementary files. Second, OTUs clustered with different levels of similarity (e.g. 97 vs 99%) are not directly comparable: if two studies are to be combined, it is necessary to perform clustering again, starting from the raw sequences. Third, taxonomic assignation is often done with different reference databases, such as GenBank, Silva or PR^2^ (Guillou et al. 2013; Quast et al. 2013). Some studies have tried to combine sets of samples from different environments (e.g. marine, freshwater and soil, Singer et al. 2021), but these efforts remain limited (for example, the Singer et al. 2021 only included 122 sampling sites). Thus, there is clearly a need to provide the protist research community with a reference database of metabarcodes which would allow the full exploration of the available sequencing data by combining existing studies across different environments.

In this paper, we introduce a database of metabarcodes (metaPR^2^) containing more than 4,150 samples originating from 41 public studies, most using the V4 region of the 18S rRNA gene. In order for the different metabarcodes to be directly comparable, we reprocessed all primary files (except those from the *Tara* Oceans expedition) with the same pipeline based on the dada2 R package (Callahan et al. 2016) and assigned the taxonomy of the resulting ASVs using PR^2^ as a reference database (Guillou et al. 2013). We have developed a web application available in several forms (website, R package, Docker container) that allow to analyse, visualize and download the data. This database will be extended in the future and should prove very useful to the protist research community.

## Material and Methods

### Dataset selection and metabarcode processing

Datasets were selected from published studies (Table 1). Raw sequence files and metadata were downloaded from NCBI SRA website (https://www.ncbi.nlm.nih.gov/Traces/study) when available or obtained directly from the investigators. Information about the study and the samples (substrate, size fraction etc…) as well as the available metadata (geographic location, depth, date, temperature etc…) were stored in three distinct tables in a custom MySQL database. For each study (except for the V9 *Tara* Oceans dataset, see below), raw sequences files were processed independently *de novo* on the Roscoff ABIMS (Analysis and Bioinformatics for Marine Science) cluster. Primer sequences were removed with *cutadapt* (Martin 2011) using the default parameters (maximum error rate = 10%). Amplicon processing was performed under the R software (R Development Core Team 2013) using the *dada2* package (Callahan et al. 2016). Read quality was visualized with the function plotQualityProfile. Reads were filtered using the function filterAndTrim, adapting parameters (truncLen, minLen, truncQ, maxEE) as a function of the overall sequence quality. Merging of the forward and reverse reads was done with the mergePairs function using the default parameters (minOverlap = 12, maxMismatch = 0). Chimeras were removed using removeBimeraDenovo with default parameters. For the *Tara* Oceans dataset, because of the very high read coverage, we did not reprocess the Illumina read files and used the sequences that had been and clustered with the *Swarm* software (Mahé et al. 2014) as detailed in de Vargas *et al*. (2015). ASVs with similar sequences from different studies were merged together and identified with a unique 10 character code which corresponds to the start of 40-character hash value of the sequence (using the R function digest::sha1). Taxonomic assignation of all ASVs, including those from *Tara* Oceans, was performed using the assignTaxonomy function from *dada2* against the PR^2^ database (Guillou et al. 2013) version 4.14 (https://pr2-database.org/). ASV sequence and taxonomy, as well as abundance in each sample, were stored in MySQL tables in the same database as the metadata (see above). In order to limit the size of the database, we only considered ASVs that corresponded to more than 100 reads in any given studies. The number of reads in each sample was normalized to 100 such that the read abundances could be expressed as % of total eukaryotic reads in some visualizations (e.g. in maps, see below). We also did not consider sequences that had an assignment bootstrap value lower than 75% at the supergroup level. Sequence processing scripts can be found in https://github.com/vaulot/Paper-2021-Vaulot-metapr2/tree/main/R_processing.

**Table 1:**
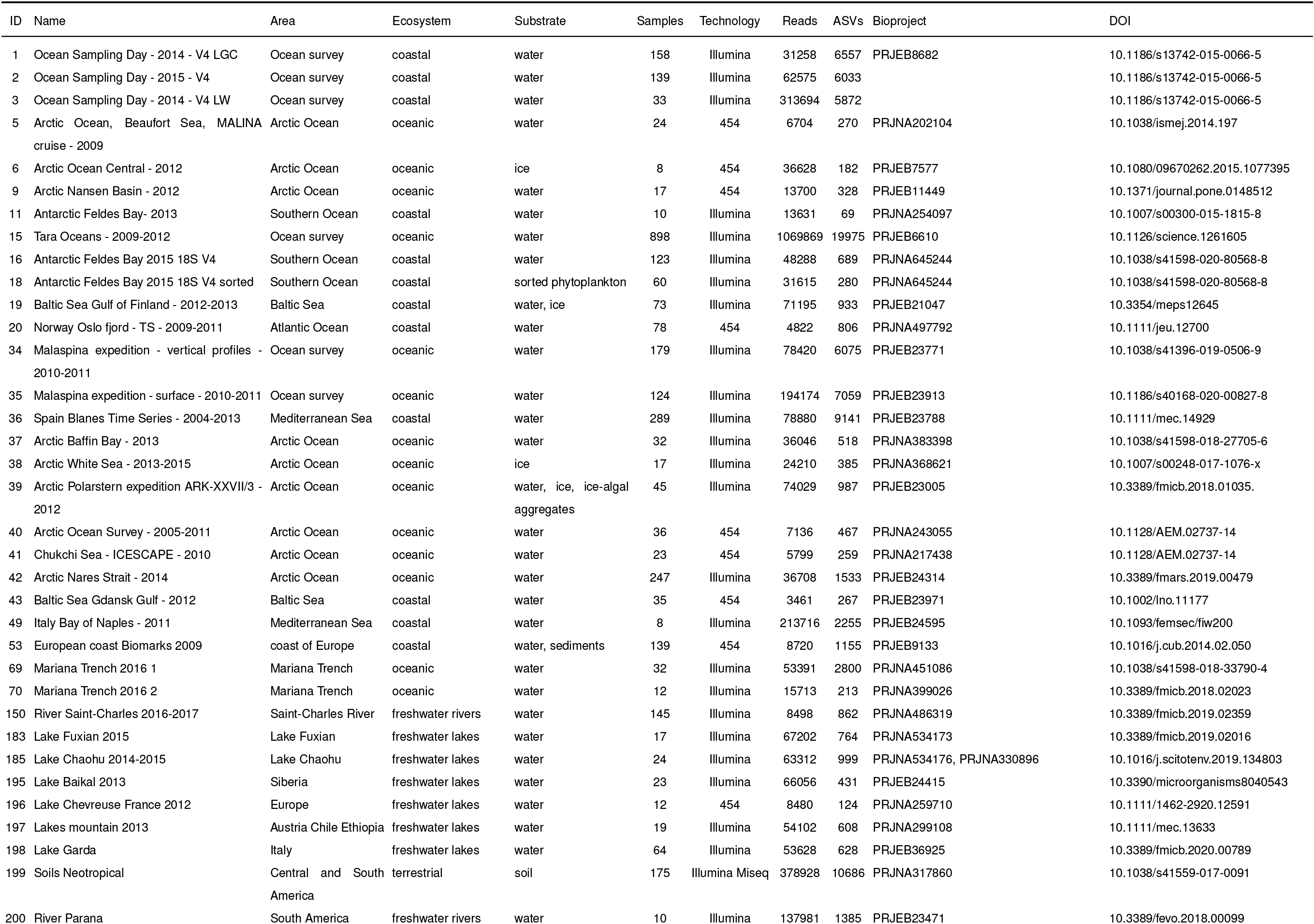
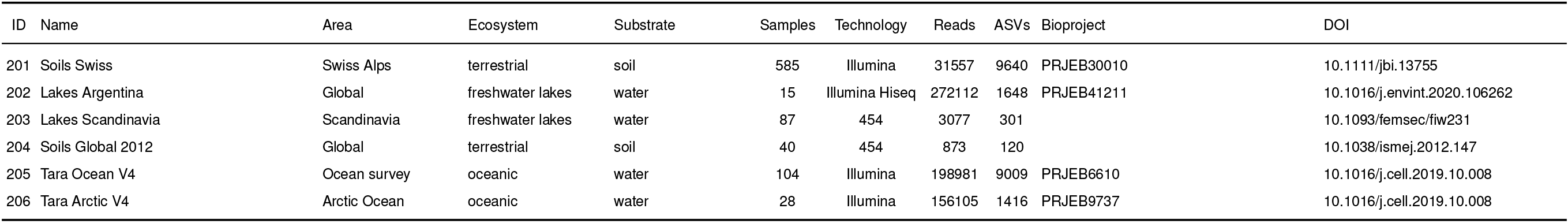
List of eukaryotic datasets and studies included in the metaPR2 databases. The column ‘Reads’ corresponds to mean number of reads per sample.

### Metabarcode analysis

Since the datasets included into metaPR^2^ used different sets of primers (see below Table S3), we clustered ASVs with 100% similarity using *vsearch* –cluster_fast option. ASVs within each cluster were merged together, using the centroid ASV as the new ASV. In order to evaluate the similarity of ASVs to existing sequences, we followed the approach of Metz et al. (2021). We compared ASVs to sequences from the PR^2^ database (Guillou et al. 2013) version 4.14 (https://pr2-database.org/) using the *vsearch* –usearch_global function with iddef = 2. The similarity information was stored in the MySQL database and then retrieved and merged with the ASV information using an R script. Alpha and beta diversity analyses were performed using the R *phyloseq* package (McMurdie and Holmes 2013).

### Ecological function

We used the table provided in Table S2 of Sommeria-Klein et al. (2021) which defines one of 4 ecological functions (phototroph, phagotroph, parasite, metazoa) to taxonomic groups (mostly at the class or division level). This table was merged with the PR^2^ taxonomy table, propagating the ecological function down to the species level. For taxonomic groups for which the paper had not defined any function, we complemented it based on general knowledge for protists (see Table S1)

### R shiny application

All post-processing was done with the R software. The data were extracted from the MySQL database using a custom script and stored in files using the R *qs* package that allows extremely fast loading of files (Travers 2021). The data are post-processed using packages *dplyr* and *tidyr*. An R shiny application was developed to interact with the database using the following R packages: *shiny, DT, shinyvalidate, shinyWidgets* and *shinycssloaders* (Sali and Attali 2020). Data are plotted using packages *ggplot2, treemapify, leaflet, leaftlet*.*minipie* and *plotly*. Alpha and beta diversity analyses are performed using the *phyloseq* package (McMurdie and Holmes 2013). The shiny application is available in 3 forms: a web-based application (https://shiny.metapr2.org), an R package (https://github.com/pr2database/metapr2-shiny) or a Docker container (https://hub.docker.com/repository/docker/vaulot/metapr2). The web interface is running on a Google Cloud Virtual Machine with a 10 Go virtual disk and 4 Go of memory. Both the R package and the Docker container can be installed on any computer.

## Results and Discussion

### Overview of metaPR^2^ datasets

Forty-one datasets are included in the first version of the metaPR^2^ database (Table 1). We selected global oceanic datasets (OSD, Malaspina, *Tara* Oceans) that have been used in numerous publications (e.g. Giner et al. 2020; Ibarbalz et al. 2019; Tragin and Vaulot 2018) as well as smaller data sets in particular from polar waters which have been little explored. Eleven out of the 41 datasets were sequenced using the 454 technology and the rest with Illumina (mostly 2×250). The vast majority of the 41 datasets used the V4 region of the 18S rRNA gene which is the most used metabarcode to date (Lopes dos Santos et al. 2021), with only two datasets representing the V9 region (*Tara* Oceans and Argentinian lakes, Table 1). The most common primer pairs used for V4 (Figure S1, Table S2 and S3) were those designed by Stoeck et al. (2010) and modified by Piredda et al. (Piredda et al. 2017). The V4 metabarcodes varied from 309 bp to 672 bp and were overlapping (Figure S1).

The metaPR^2^ database contains more than 4,150 samples (Figure 1). These samples originate from three major ecosystems: marine, freshwater and terrestrial (mostly soil substrate) (Figure 2). Among water samples, different size fractions from pico (0.2-3 *μ*m) to meso (100-1000 *μ*m) are represented with the majority corresponding to the pico and total fractions (Figure 2). Most aquatic samples correspond to surface or euphotic layer. Location data (longitude, latitude) are available for all samples but other metadata, e.g. temperature or salinity, may be missing for some samples (Figure S2).

**Figure 1:**
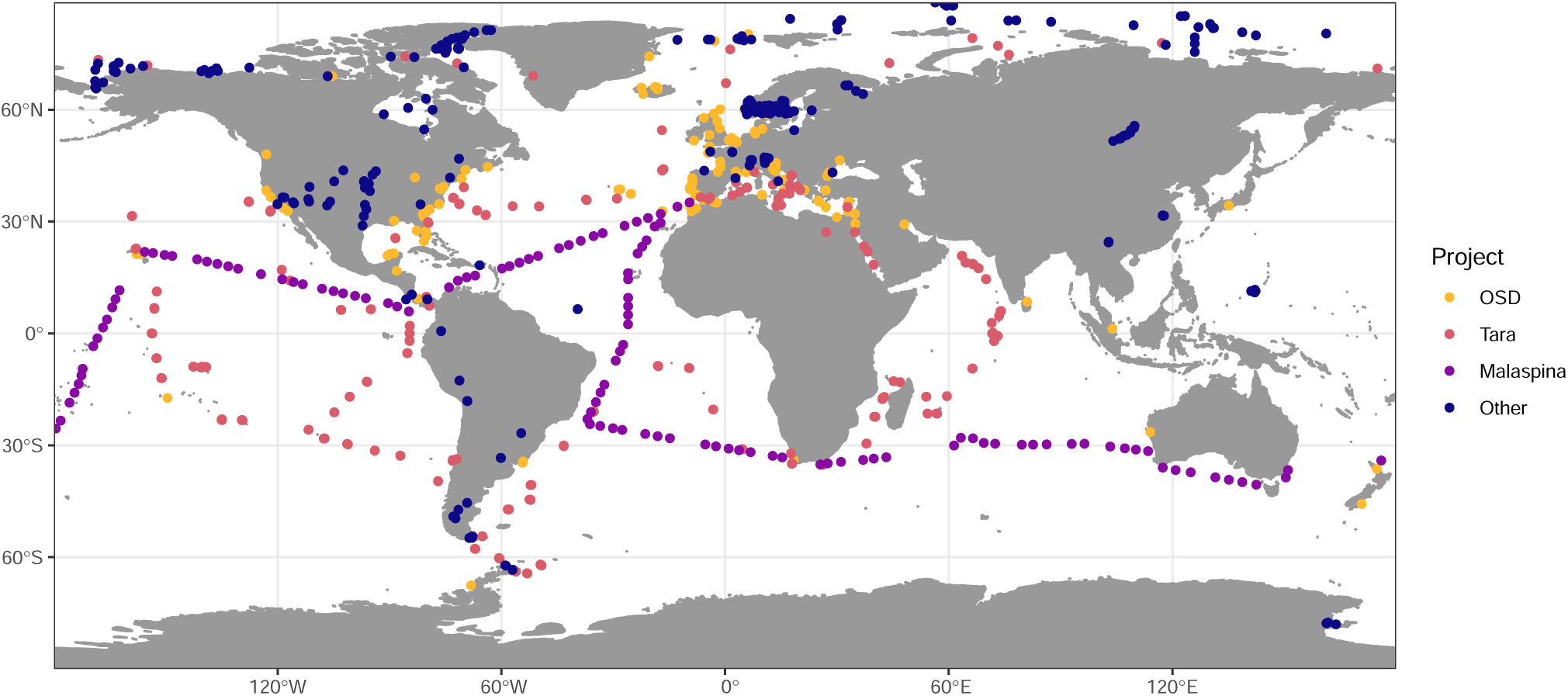
Map of stations included in the metaPR^2^ database.

**Figure 2:**
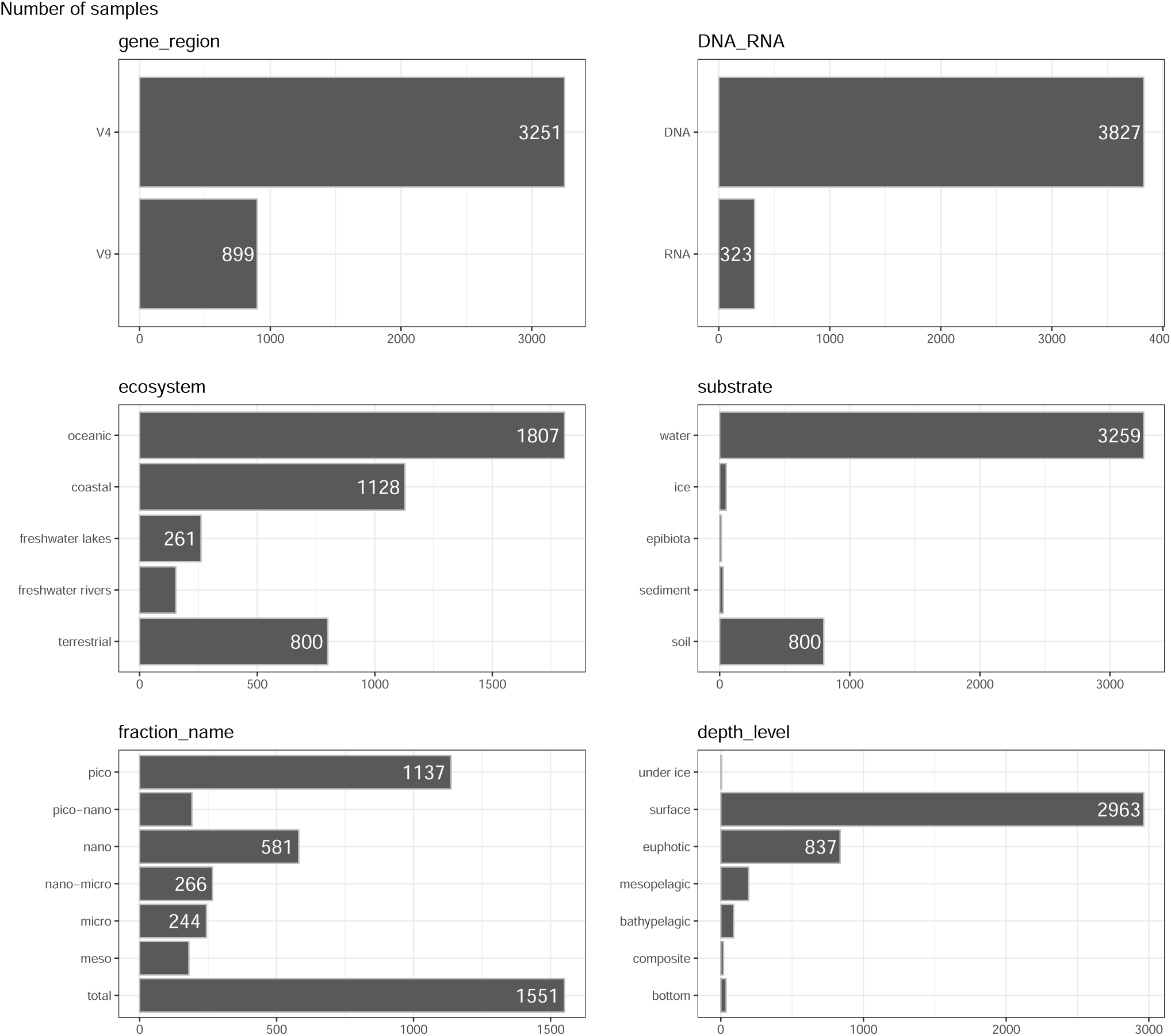
Distribution of samples by gene region, DNA or RNA, ecosystem, substrate, fraction name and depth level.

The number of samples per dataset is quite heterogeneous ranging from less than 10 to almost 900 for *Tara* Oceans (Table 1). The total number of reads analysed is almost 900 million for V9 and above 220 million for V4. The number of reads per dataset is also highly variable ranging from about 3,000 in the older studies sequenced by 454 technology to more than 1 million for Tara V9 (Table 1), which explains why overall there are more reads for V9 than V4 despite only 2 datasets using V9. The total number of ASVs was about 79,000. The number of ASVs in a given study ranges from less than 100 to more than 14,000 depending on both the number of samples and the depth of sequencing (Table 1). Since different studies have used different primer sets, it is necessary to cluster ASVs with 100% similarity in their shared region, leading to slight reduction of the total number of ASVs from 79,000 to 70,000 once clustered. In general, sequences included in a given cluster were widely overlapping, although a few bases could be different outside the overlap region, pointing to some microdiversity within these clusters (Figure S3). All results presented below used the clustered ASVs that we call cASVs.

### Protist composition

Overall, the database is dominated by Opisthokonta (Metazoa and Fungi) and Alveolata (Dinoflagellata) (Figure S4). In what follows, we decided to focus on protists and on the V4 region. The focus on protists is justified because the sampling strategy of most datasets was optimal for microbial eukaryotes. DNA from those three divisions not included in protists (metazoa, plants and fungi) were probably unevenly sampled, e.g. plant seeds in soils, multicellular organism, larval stages of metazoa in water environments. The focus on the V4 datasets that contain almost 3,000 samples and 850 sites is due to that fact that the data for the V9 region are dominated by the *Tara* Oceans dataset, which has been extensively analysed previously (e.g., De Vargas et al. 2015).

Protist sequences represent more than 40,000 ASVs (33,000 cASVs once clustered). In terms of reads and cASVs, the database is dominated by Alveolata, followed by Stramenopiles, Hacrobia, Archaeplastida and Rhizaria (Figure 3). Based on number of cASVs, Rhizaria despite their lower read abundance come just after the Stramenopiles. Such large number of Rhizaria unique sequences compared to read numbers has been observed before, possibly linked to higher error rates in regions of the RNA molecule that form secondary structures (2011). The most abundant cASVs (Figure 4A) belong to dinoflagellates (*Gyrodinium*), diatoms (*Minidiscus, Porosira, Fragilariopsis*), cryptophytes (*Geminigera, Cryptomonas*), haptophytes (*Phaeocystis*) and green algae (*Bathycoccus, Micromonas*). The most abundant cASVs are often also the most frequently occurring (Figure 4B and C), although for example the marine picoplanktonic genus *Florenciella* is quite frequent despite being not one of the most abundant. In contrast, the abundant small diatom *Minidiscus* cASV is not present among the 30 most frequent cASVs. The difference in reads abundance and cASV frequency among these two marine phytoplanktonic genera might be a reflection of their coastal-oceanic distribution, which can be easily observed with the online platform of metaPR^2^. *Florenciella* is truly a ubiquitous genus, found in both coastal and oceanic samples, although often in low abundance. In contrast, the nanoplanktonic diatom *Minidiscus* is mostly found in coastal environments or continental platforms, where it can form sporadic blooms (Leblanc et al. 2018).

**Figure 3:**
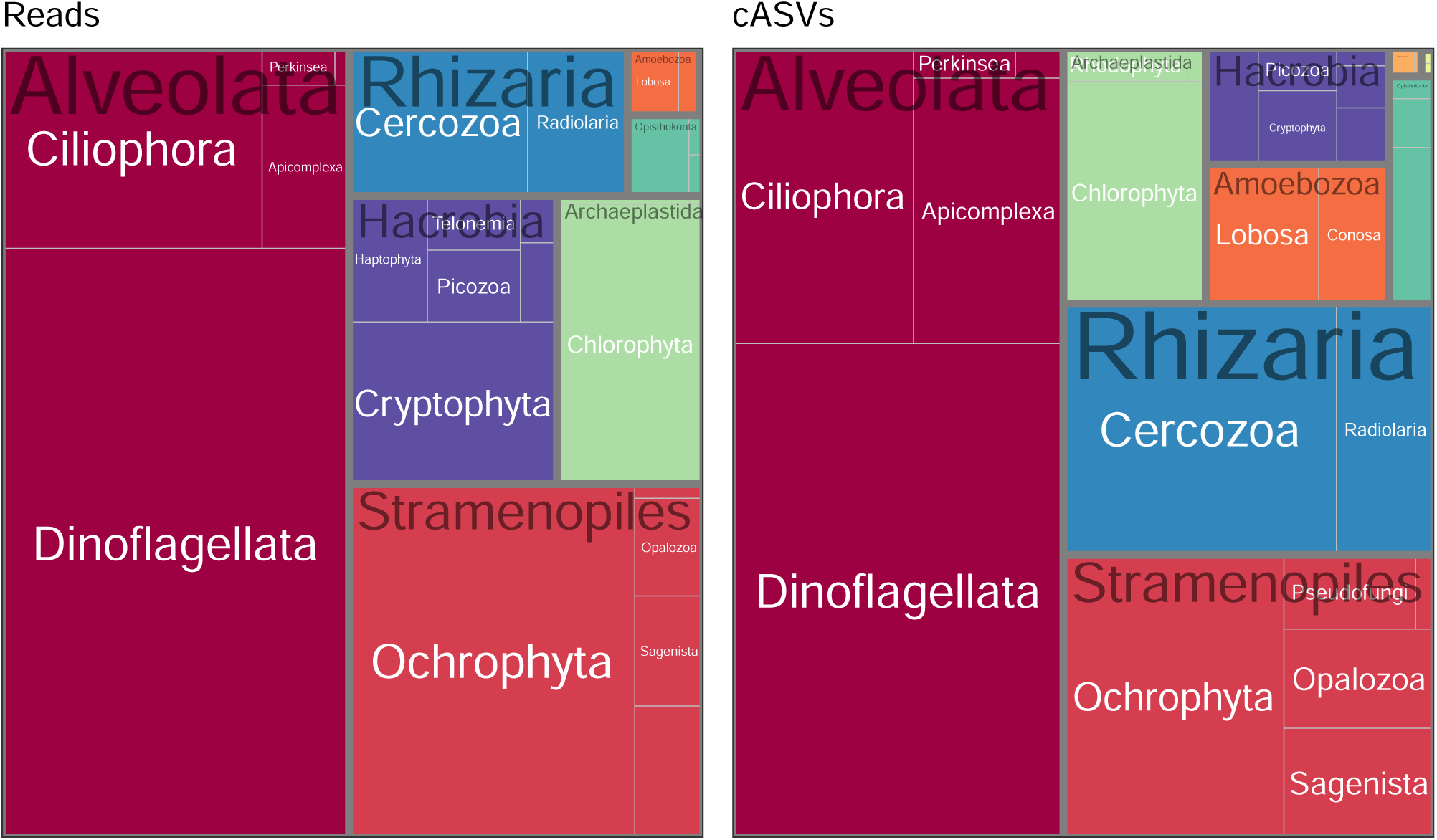
Treemaps of most abundant protist taxa (supergroup and division) for V4 datasets based on number of reads after normalization (left) or number of cASVs (right).

**Figure 4:**
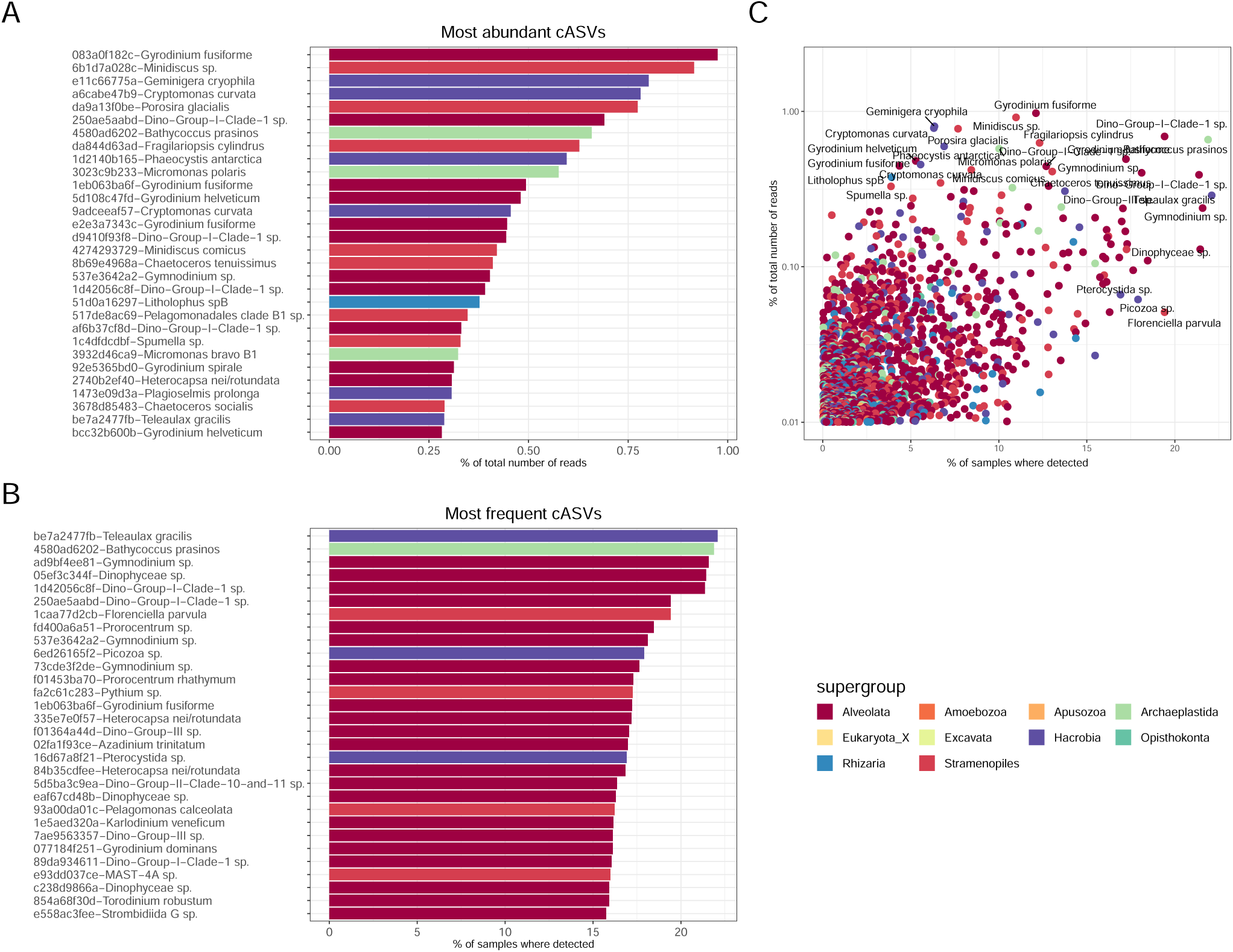
Protist V4 cASVs. Most abundant cASVs (after normalisation per sample). B. Most frequent cASVs. C. Relationships between cASV frequency and abundance. Each cASV is coded by a 10-letter string representing the start of the 40-character hash value of the sequence (see Material and Methods).

Comparing the metaPR^2^ metabarcodes to reference databases such as PR^2^, reveals that there are very few novel metabarcodes for supergroups such as Hacrobia and Archaeplastida that contain many photo-synthetic taxa. In contrast, for supergroups that contain mostly heterotrophic organisms, and in particular Amoebozoa, the median similarity of metabarcodes to any reference sequence is below 90% (Figure 5A) suggesting the existence of a lot of unknown taxa. A similar observation was recently done for a restricted set of samples from a river floodplain in Argentina (Metz et al. 2021).

**Figure 5:**
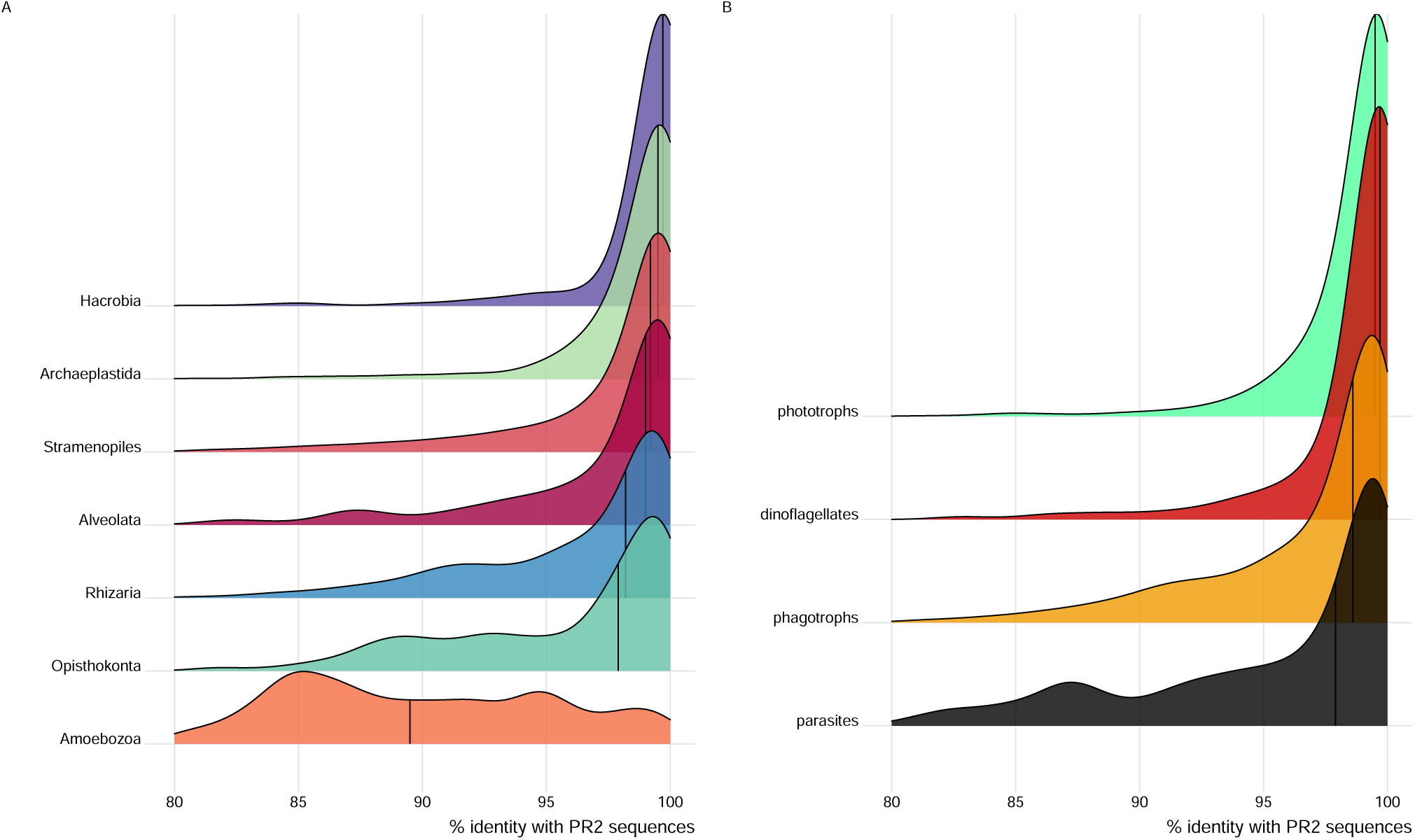
Protist V4 cASVs. Similarity of cASVs to sequences for the sequences from the PR^2^ database as a function of supergroup (A) and of the ecological function (B).

### Global trends across environments

The metaPR^2^ database corroborates some trends that have been observed in papers with much fewer samples. Singer et al. (2021) examined patterns of diversity across marine, freshwater and terrestrial (soil) ecosystems based on 122 samples. Using the metaPR^2^ database which contain 23 times more samples we are able to establish clear differences across 5 types of ecosystems: marine, coastal, freshwater lakes and rivers and terrestrial (soils). In terrestrial environments, Hacrobia are almost completely absent while Amoebozoa are present but in contrast absent in all the others environments (Figure 6A). If we use the ecological function, as defined for each major taxonomic group by Sommeria-Klein et al. (2021), the five environments clearly differ by the abundance of parasites, small number of phototrophs and absence of dinoflagellates in soils. While parasites are abundant in soils, they are not as abundant in freshwater and increase from coastal to oceanic waters (Figure 6B). In terms of diversity, using the Shannon index as an indicator, terrestrial ecosystems are most diverse, followed by rivers, oceanic, coastal with lakes the less diverse in agreement with previous analyses (Singer et al. 2021), these differences being all significant (Figure S6). Most cASVs are restricted to a single type of ecosystem with less than 2% (620 out of 33235) common to two or more if we consider coastal and oceanic ecosystems together (Figure 7). The highest number of cASVs corresponds to marine ecosystems (coastal and oceanic), followed by terrestrial and freshwater. Interestingly, both coastal and oceanic have a large number of specific cASVs with roughly 1/3 purely oceanic, 1/3 purely coastal and 1/3 common. It is also striking that there are very few cASVs common between freshwater rivers and lakes (just above 7%). In terms of novelty, i.e. of cASVs with low similarity to known sequences, terrestrial ecosystems are the least known with a median similarity below 95% followed by rivers, lakes, coastal and pelagic ecosystems (Figure 5B). In some way, this reflects the fact that soil protists have only been recently investigated (Geisen et al. 2018). A comparison between the communities structures from these different ecosystems using NMDS (Figure 8) reveals a clear gradient from terrestrial ecosystems, through rivers and lakes, towards coastal and then oceanic systems. Interestingly, river communities are the closest to soil communities, as they are probably enriched in terrestrial protists through soil drainage.

**Figure 6:**
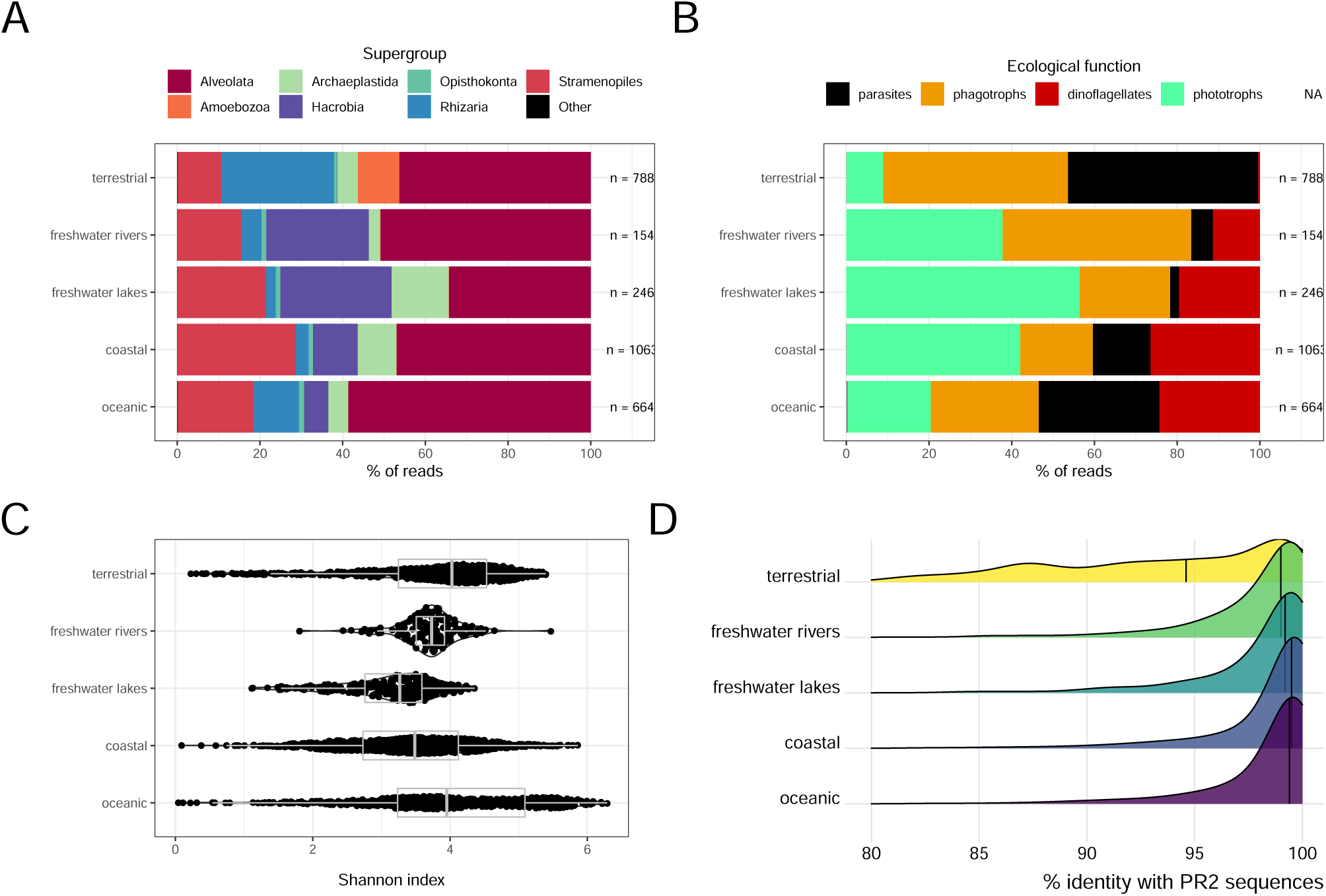
Protist V4 cASVs. Composition as a function of the environment based on taxonomy (A) or on ecological function (B) and Shannon index (C). Similarity of cASVs to sequences for the sequences from the PR^2^ database as a function of the environment (D).

**Figure 7:**
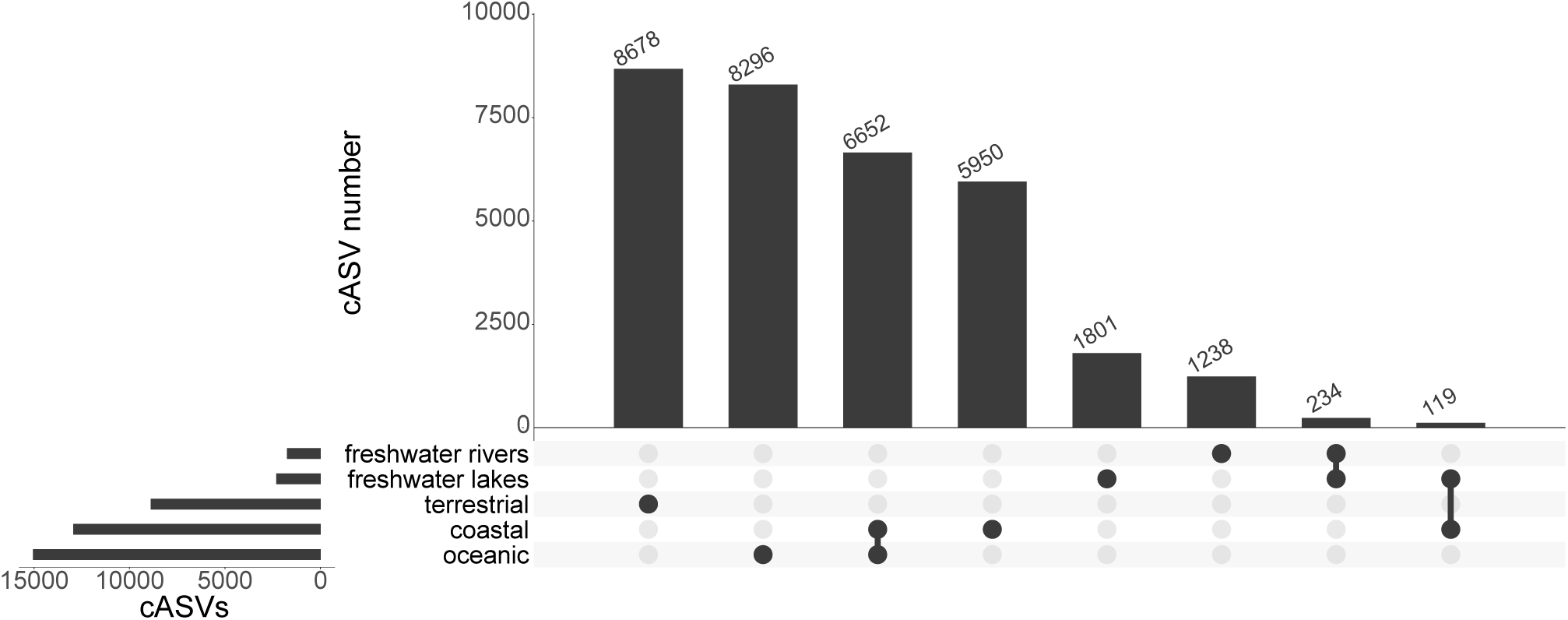
Protist V4 cASVs found on one or more environments (so-called “upset” plot).

**Figure 8:**
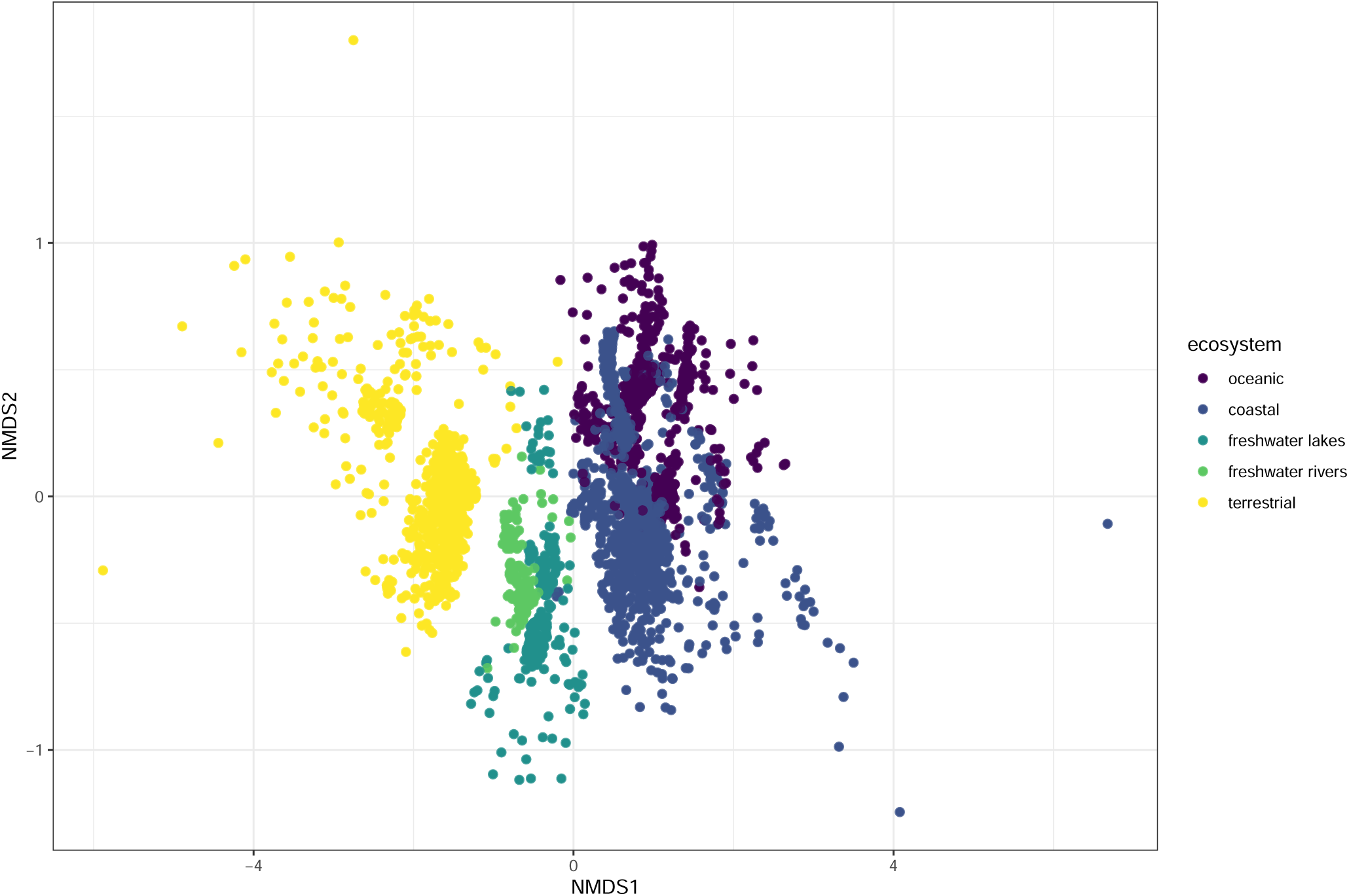
Protist V4 cASVs. NMDS analysis. Colour correspond to sample environment.

### R Shiny application

With a database of such size and complexity, it is necessary to create tools that allow in a first step to explore the database and then to download the data of interest (e.g. for a specific taxonomic group or environment). For this purpose, we developed an R Shiny application (Figure 9). R Shiny is an open source tool that offers numerous advantage to develop web-based applications in comparison to coding directly under languages such as JavaScript or PHP. It offers predefined components allowing the user to interact with the data (User Interface), while the Server component performs the necessary computations (e.g. filtering, summarizing the data etc…)in the background. Moreover, a Shiny application can be easily deployed on a server using open source tools such as Shiny Server, be packaged in a Docker container that can be downloaded on a personal computer and run locally or delivered as an R package.

**Figure 9:**
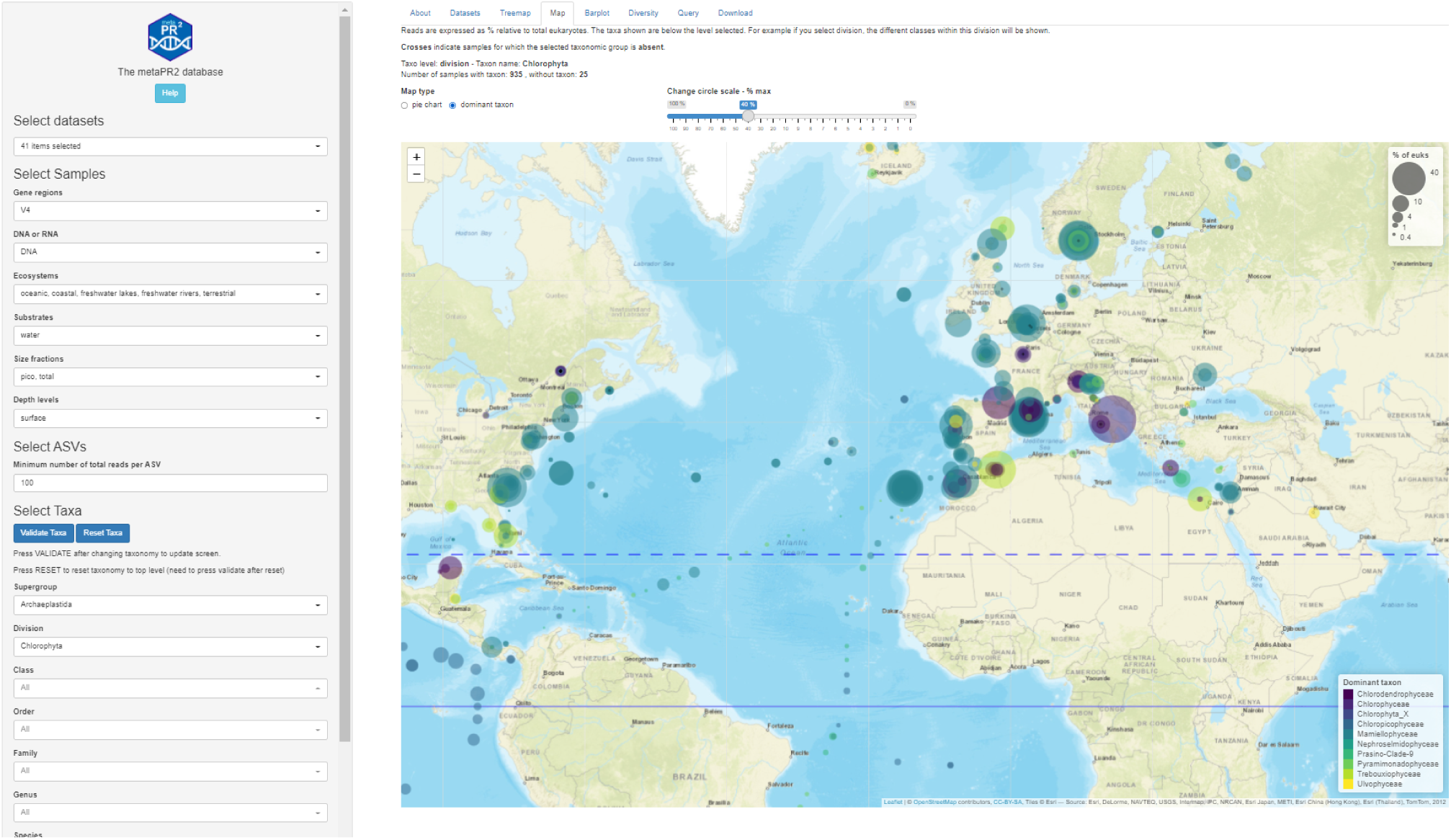
The metaPR^2^ shiny application available at https://shiny.metapr2.org.

The metaPR^2^ Shiny application is structured in a number of panels, each dedicated to one type of analysis (e.g. map, diversity). It is possible to select/deselect specific datasets or groups of datasets, such as all oceanic datasets (Figure S8). Selection can also be done based on sample characteristics such as whether samples come from DNA or RNA, the ecosystem, the type of substrate (e.g. ice, water, soil), the size fraction and the depth level (Figure S9). It is possible through reactive menus to navigate the taxonomy tree down to the cASV level (below the species) that potentially corresponding to cryptic or subspecies. cASVs can be filtered based on the number of reads found for this cASV in the whole database (between 100 and 10,000). The number of total reads for a given taxonomic level can be visualized in a treemap (Figure S10). For this representation, number of reads are normalized to 100 for each sample. The distribution of any taxon can be visualized on a map (Figure S11). Two visualization modes are proposed for maps: either a pie chart at each station with a fraction of the different taxa immediately below the level selected (for example species, if genus is the level selected) or, alternatively, a colour circle indicating the dominant taxon immediately below the level selected (for example the dominant species in the previous example). The size of the circles is proportional to the percent of reads of the taxon selected relative to the total number of eukaryotic reads. The size of the circles can be adjusted for taxa in low abundance. Another representation is in the form of barplot (Figure S12), where the x-axis represents the fraction of reads per taxon while the y-axis represents one of the variable from the metadata (depth level, temperature). For continuous variable, bins are created. This panel can also be used for time series with different levels of aggregation (year, month, day). Alpha and beta diversity (Figure S13) can be computed for a limited number of samples (1,000 maximum). It is possible to query the whole set of cASV using a BLAST like query and to map the resulting cASVs (Figure S14). Finally, it is possible to download datasets and samples metadata as well as the cASV and the read abundance for the datasets, samples and taxa selected (Figure S15).

The metaPR^2^ shiny application besides being very useful for research can also be used for pedagogical purposes. MetaPR^2^ can be used as a tool by professors and instructors in the field of microbial ecology. In the framework of the undergraduate course ES2304 - Microbes in Natural Systems at Nanyang Technological University (Singapore), the application was used to investigate the biogeography of several groups of phytoplankton (diatoms, bolidophytes, dinoflagellates, green algae) by groups of 4 students in a flipped-classroom model. Each group had to do some research on the genus it was assigned and then to analyse the distribution and diversity of key species, answering questions such as whether some species had ubiquitous distributions or controlled by latitude or temperature and whether some species appeared to contain different genotypes as reflected by the presence of several cASVs. In order to make their analysis less daunting, they only analysed the OSD, Malaspina and Tara V4 datasets. Despite the fact that they had only one week to discover the interface and produce their analyses, this hands-on experience resulted in very positive feedbacks by the students, especially regarding using the platform to look at “real-world research data”.

### Perspectives

As its sister database PR^2^ which is revised every 6-12 months with the addition of novel sequences as well update in taxonomy, the metaPR^2^ will evolve with time to include more datasets and more samples, in particular from ecosystems (e.g. extreme environments), regions (e.g. tropical and southern latitudes) and substrate (microbiomes) that are still little represented. We have tabulated more than 280 metabarcoding studies of protist diversity, for most of which data are available from GenBank SRA. These data will be processed and incorporated into the database with probably yearly releases. The taxonomy of metaPR^2^ will evolve in parallel to that of PR^2^ and we will add other functional and phenotypic traits (e.g. size, mixotrophy type) as there is clear tendency to use this approach more widely for protists (Schneider et al. 2020). We will also develop novel functionalities for the R shiny application and package, for example heatmaps and phylogenetic analyses. This will constitute a very rich resource that will help to compare eukaryotic communities across habitats.

## Data availability

Source code for the Shiny server is available as an R package from GitHub (https://github.com/pr2database/metapr2-shiny, DOI: 10.5281/zenodo.5992354). Source code for this paper is available from GitHub (https://github.com/vaulot/Paper-2021-Vaulot-metapr2). Source code for sequence processing is available from GitHub https://github.com/vaulot/Paper-2021-Vaulot-metapr2/tree/main/R_processing.

## Acknowledgements

We thank Javier de Campo, Catherine Ribeiro, Ana Maria Cabello for their shiok suggestions on the shiny application. We thank the ABIMS platform of the FR2424 (CNRS, Sorbonne Université) for bioinformatics resources. The work of MJ and CB in the Burki lab was supported by a grant from Science for Life Laboratory available to F. Burki.

## Author contributions statement

DV conceived the study. DV, AL, DO, BT, CB scanned the literature and metadata. DV, DO, BT, MJ, CB collected and compiled metadata from the different datasets. DV developed the database structure, the analysis scripts and the R shiny application. DV performed the metabarcode analyses. CS compiled the functional trait information. DV and AL wrote the first draft of the paper, and all co-authors edited and approved the final version.

## Supplementary Material

**Table S1:**
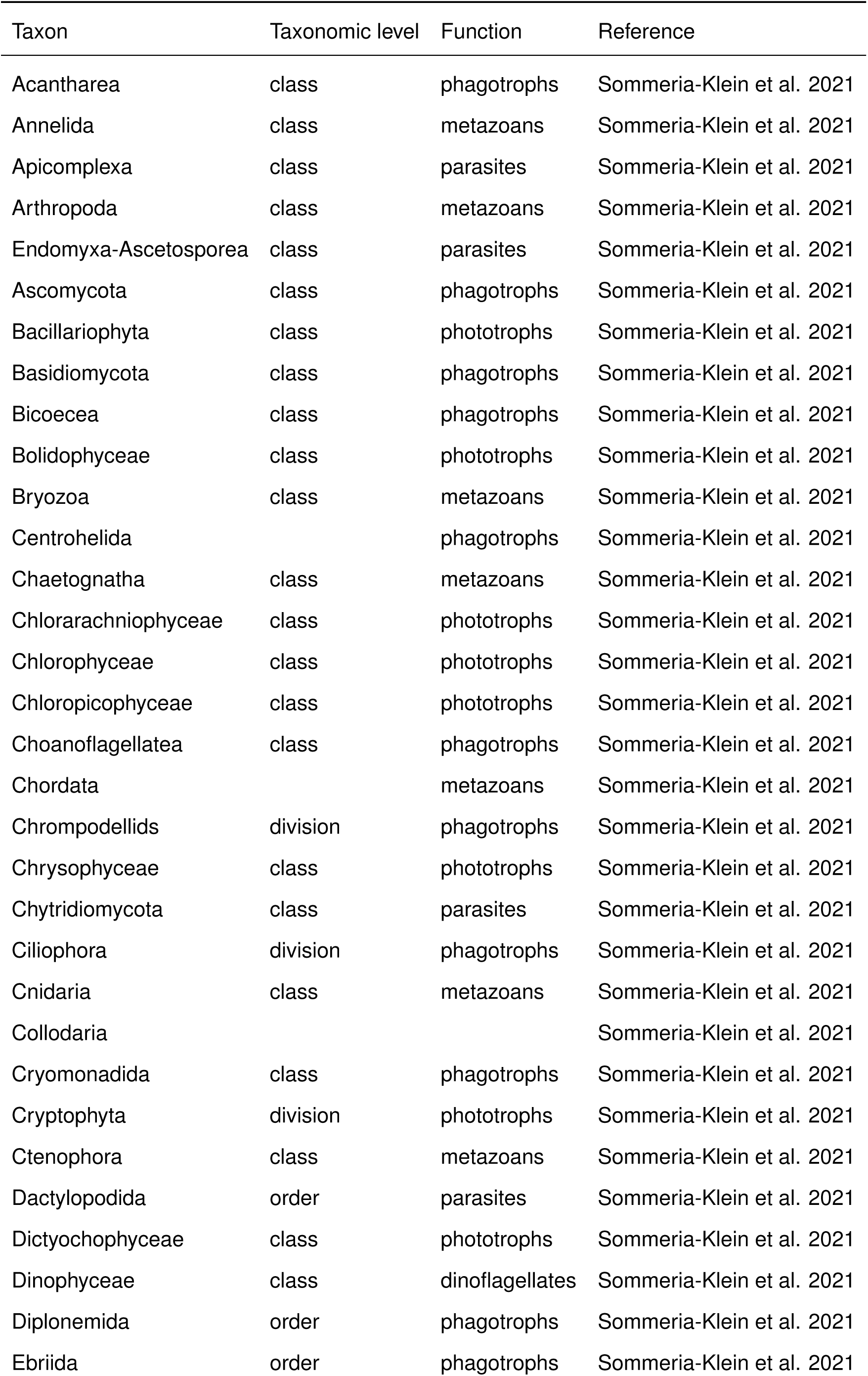
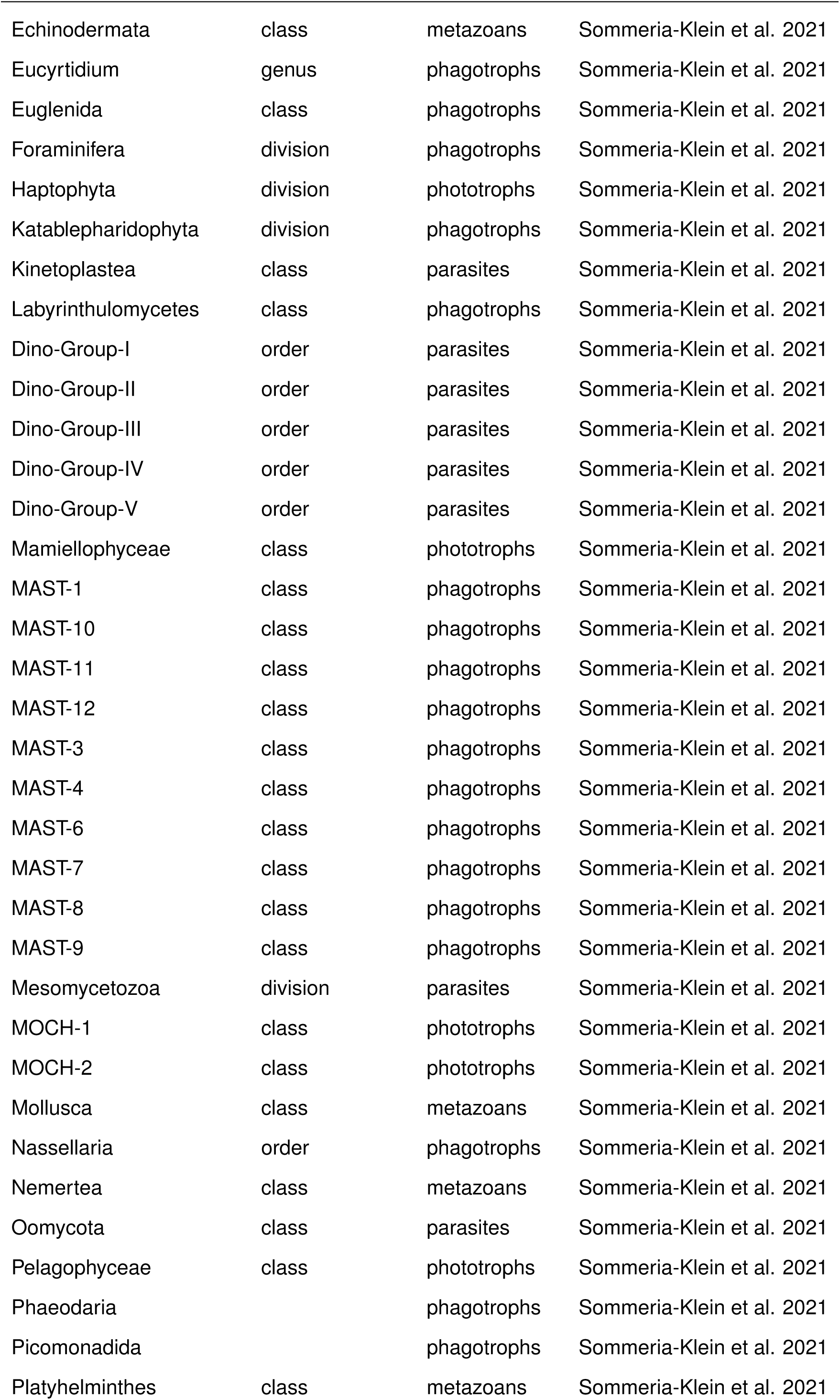
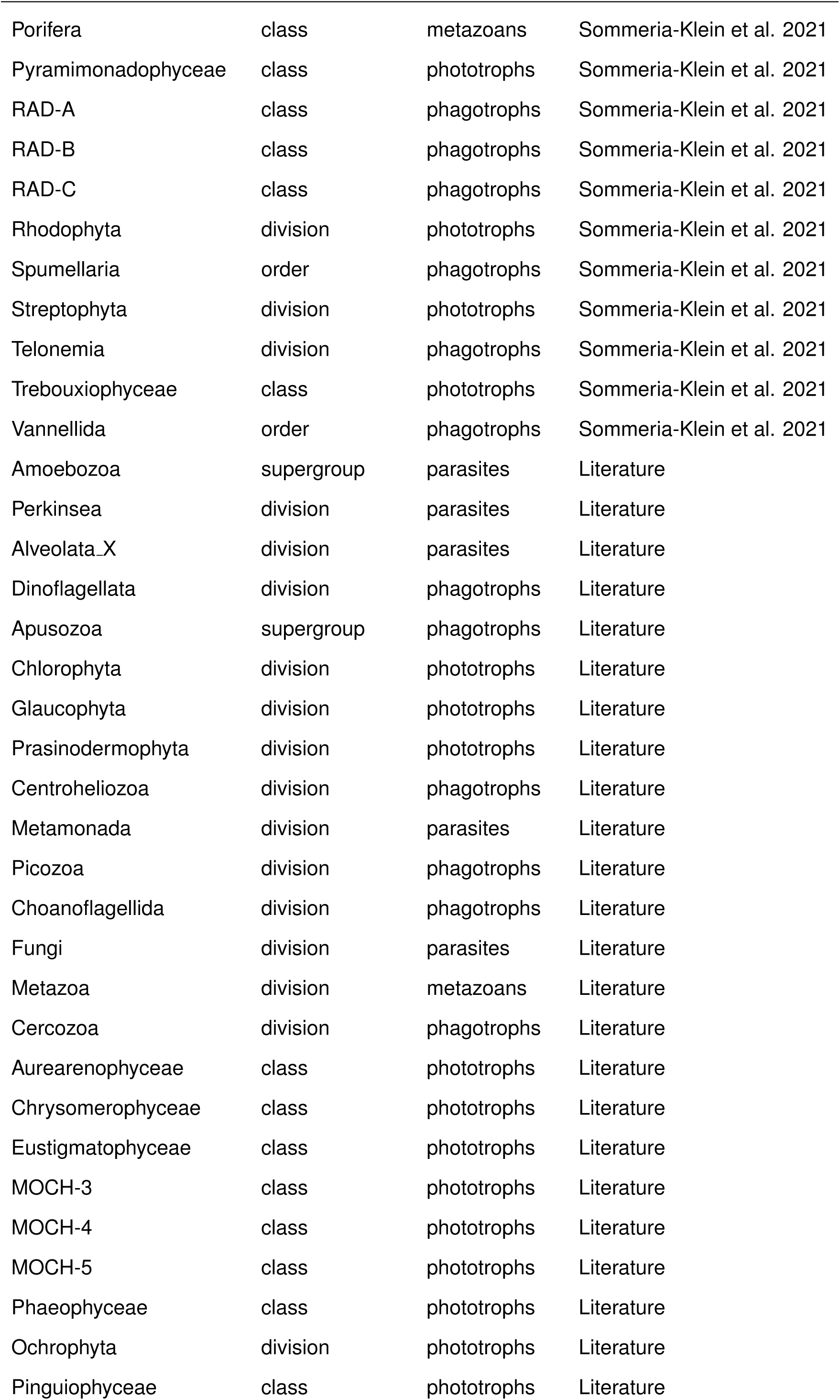
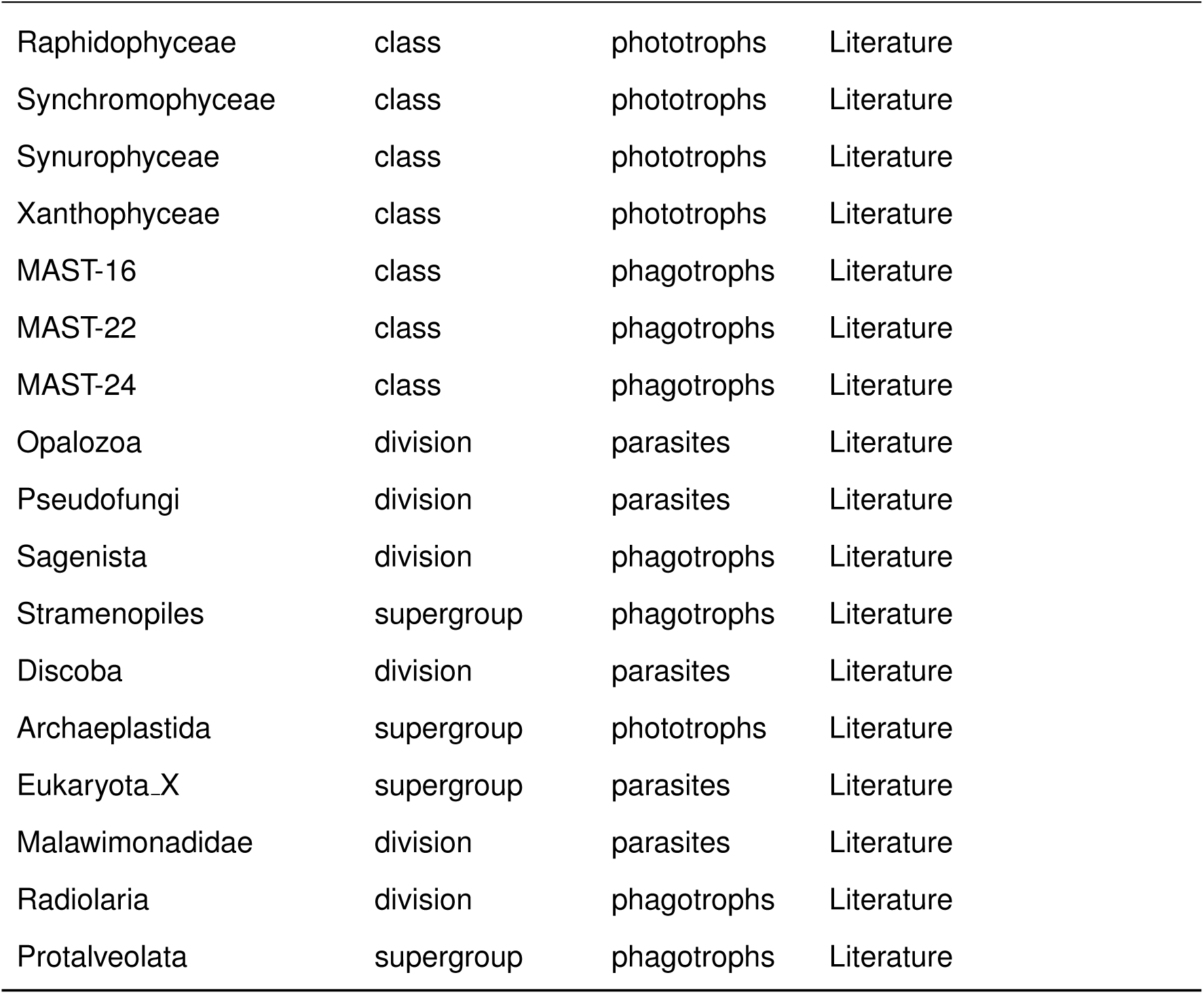
Ecological function of taxa according to Table S2 of Sommeria-Klein et al. (2021). Taxa present in the PR2 database for which ecological function was not present in Table S2 were assigned an ecological function based on the literature. Ecological function was propagated to all taxa below the taxon for which it was defined using an R script.

**Table S2:**
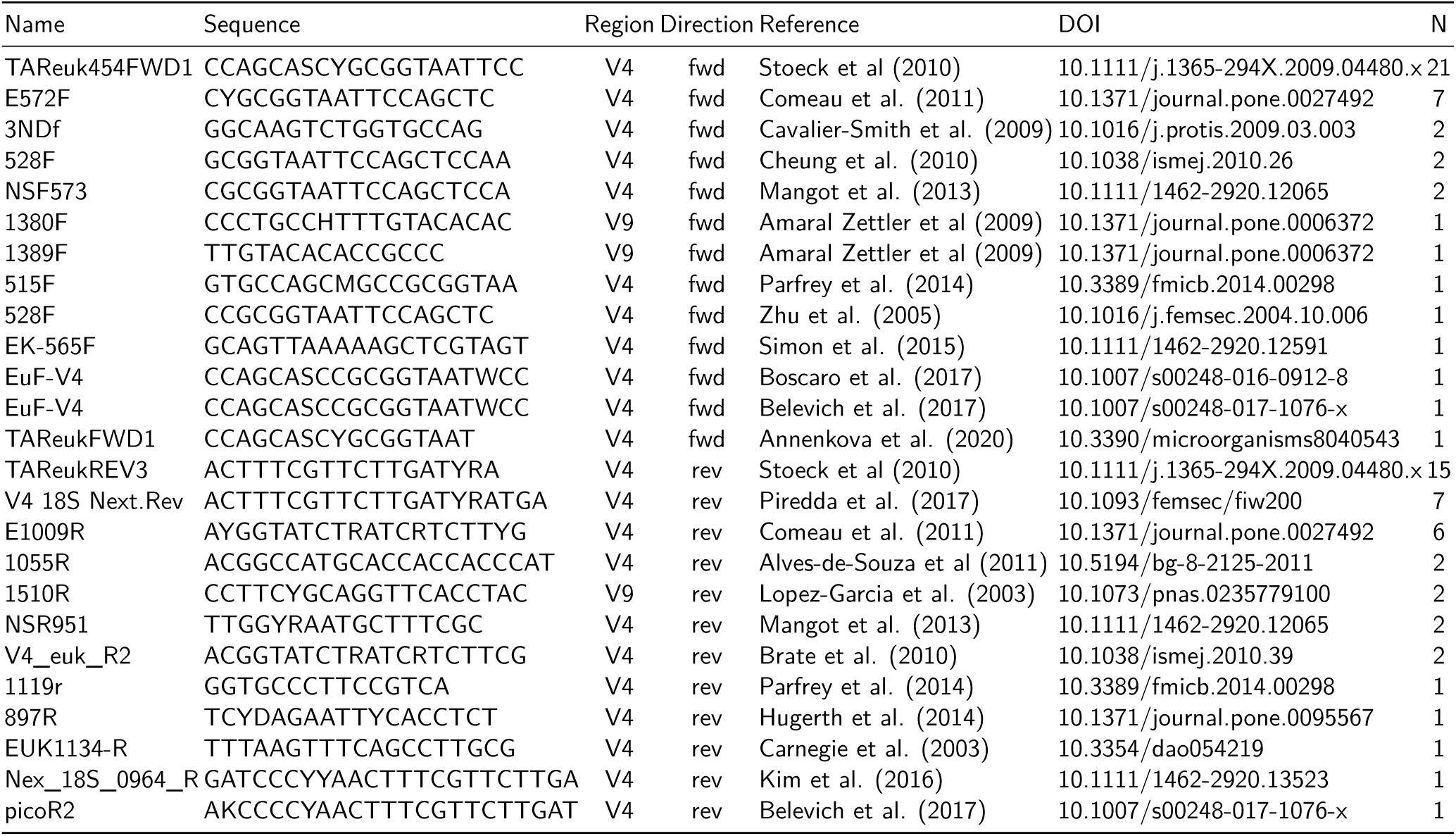
Eukaryotic 18S rRNA primers used for metaPR2 datasets with the number of datasets (N) where used (Table 1).

**Table S3:** 18S rRNA primer sets used for metaPR2 datasets with the number of datasets (N) where used (Table 1). Refer to Table S2 for sequence and reference of primers.

| Primer fwd | Primer rev | Region | N |
| --- | --- | --- | --- |
| TAReuk454FWD1 | TAReukREV3 | V4 | 14 |
| TAReuk454FWD1 | V4 18S Next.Rev | V4 | 7 |
| E572F | E1009R | V4 | 6 |
| 3NDF | V4_euk_R2 | V4 | 2 |
| 528F | 1055R | V4 | 2 |
| NSF573 | NSR951 | V4 | 2 |
| 1380F | 1510R | V9 | 1 |
| 1389F | 1510R | V9 | 1 |
| E572F | 897R | V4 | 1 |
| EK-565F | UNonMet | V4 | 1 |
| EuF-V4 | picoR2 | V4 | 1 |
| F515 | R119 | V4 | 1 |
| Nex_18S_0587_F | Nex_18S_0964_R | V4 | 1 |
| TAReukFWD1 | TAReukREV3 | V4 | 1 |

**Figure S1:**
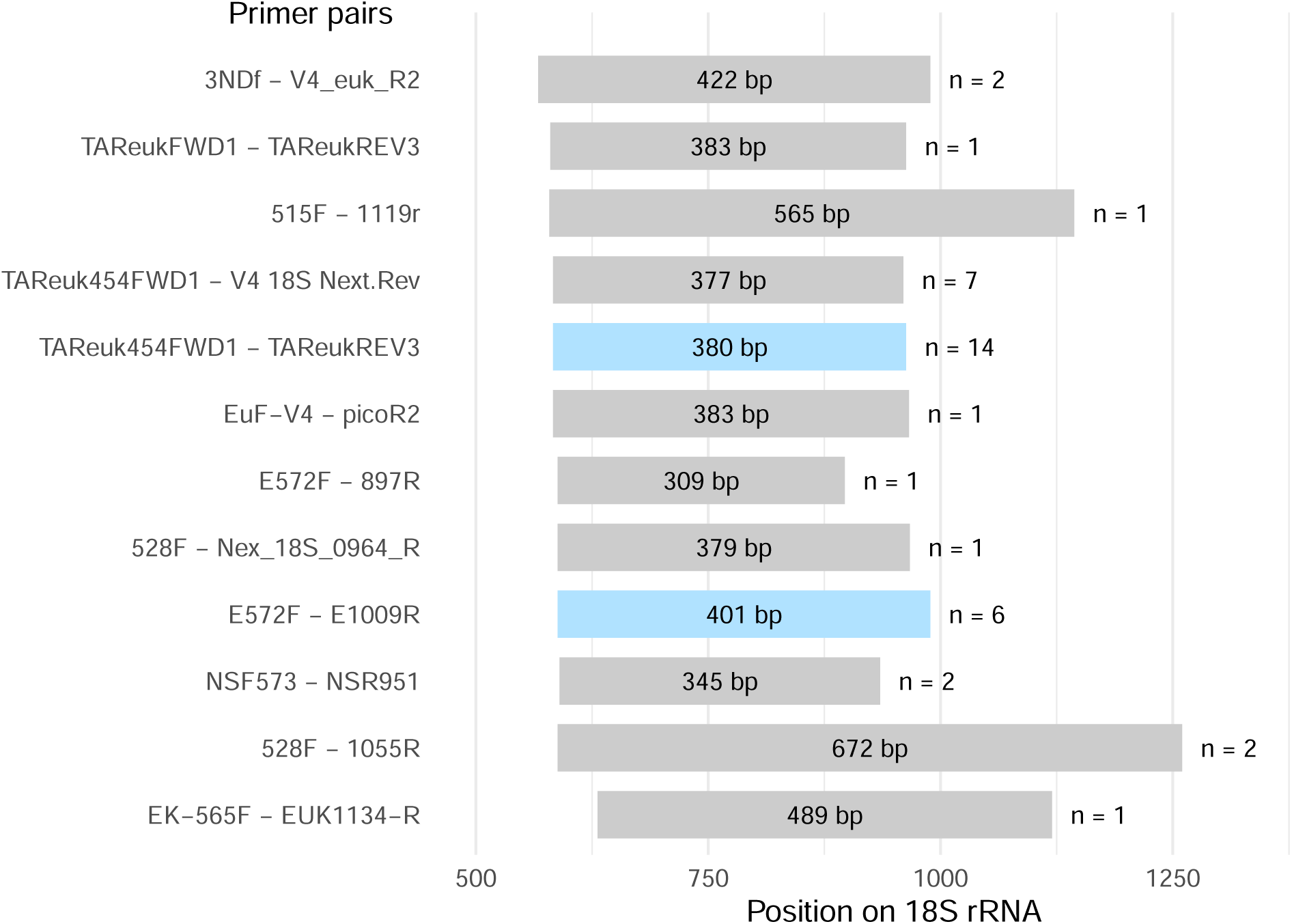
Amplicon size and position on the 18S rRNA gene (yeast), with the number of datasets for each V4 primer pair on the right side.

**Figure S2:**
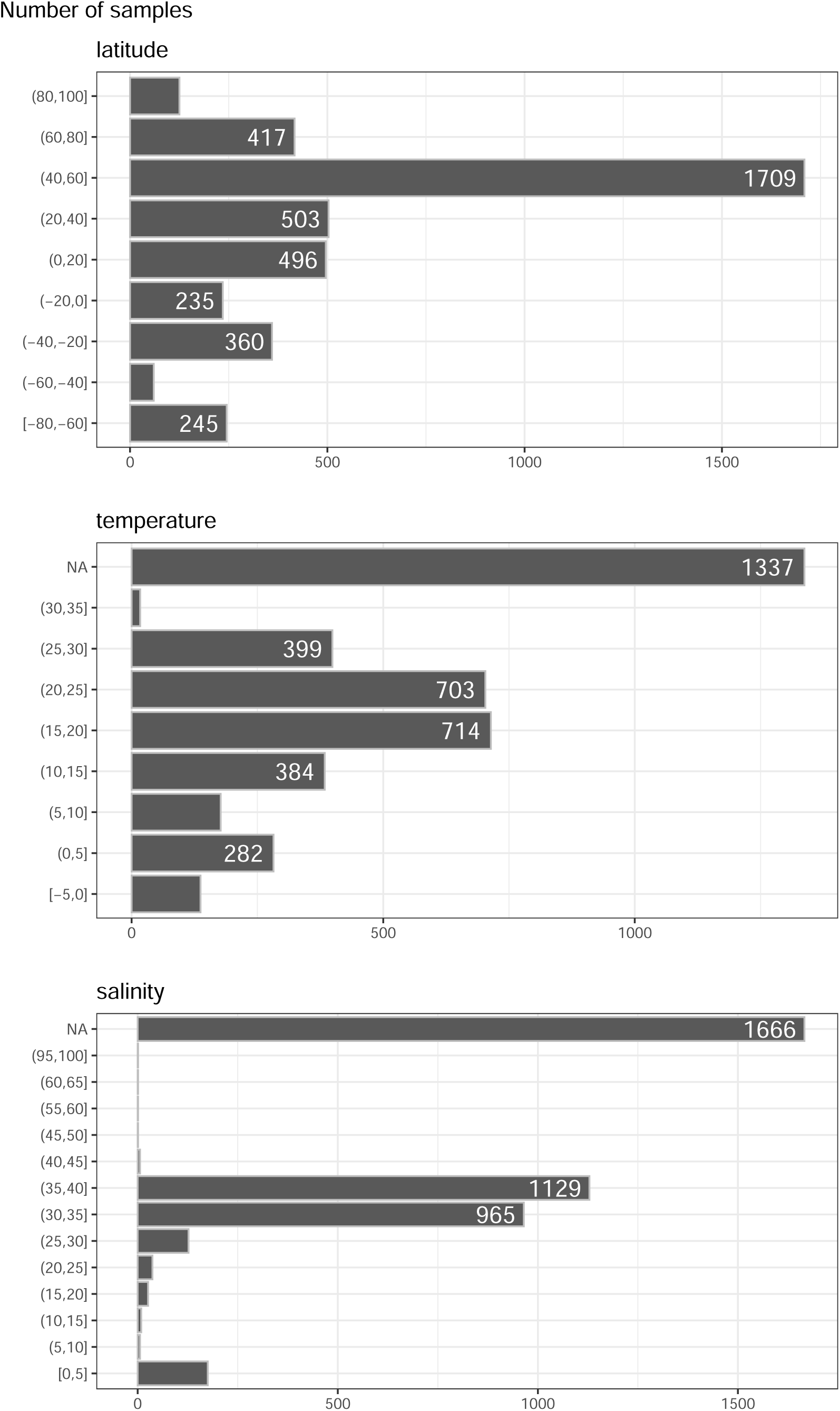
Distribution of samples by latitud, temperature and salinity ranges. NA corresponds to samples for which the data are not available.

**Figure S3:**
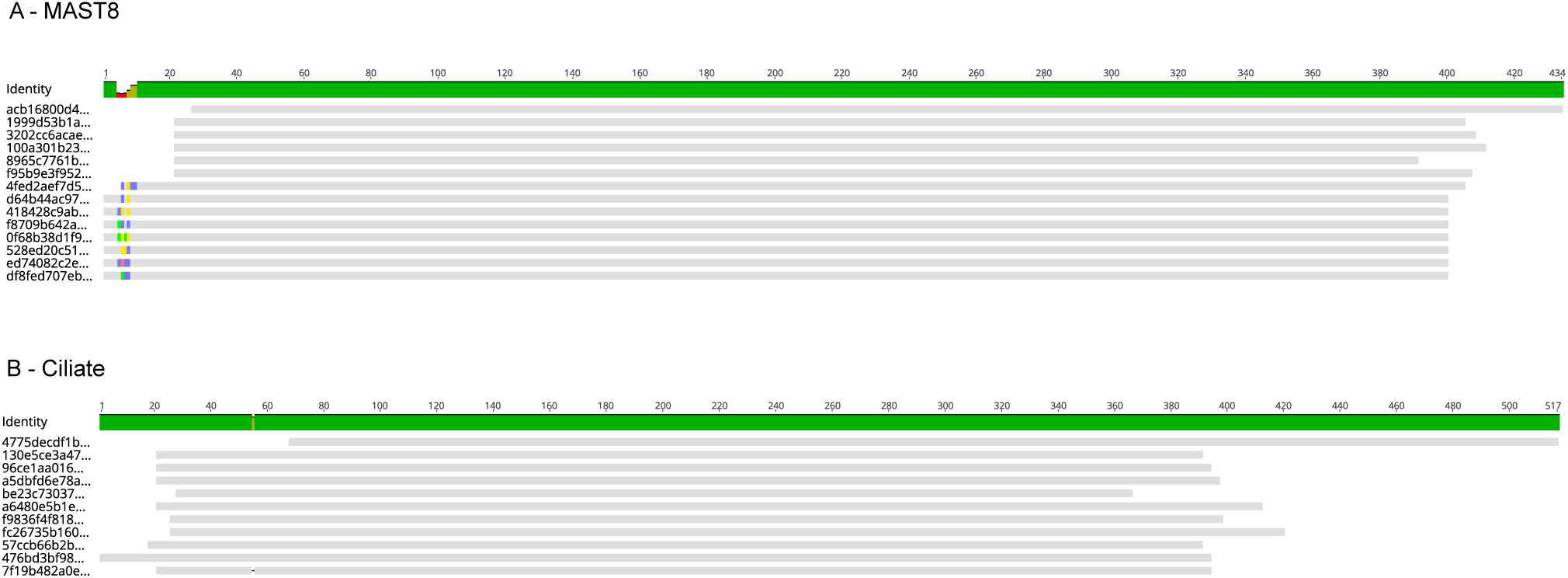
Two examples of V4 sequence clusters ASV (cASV) for Stramenopiles MAST 8 (A) and ciliates (B).

**Figure S4:**
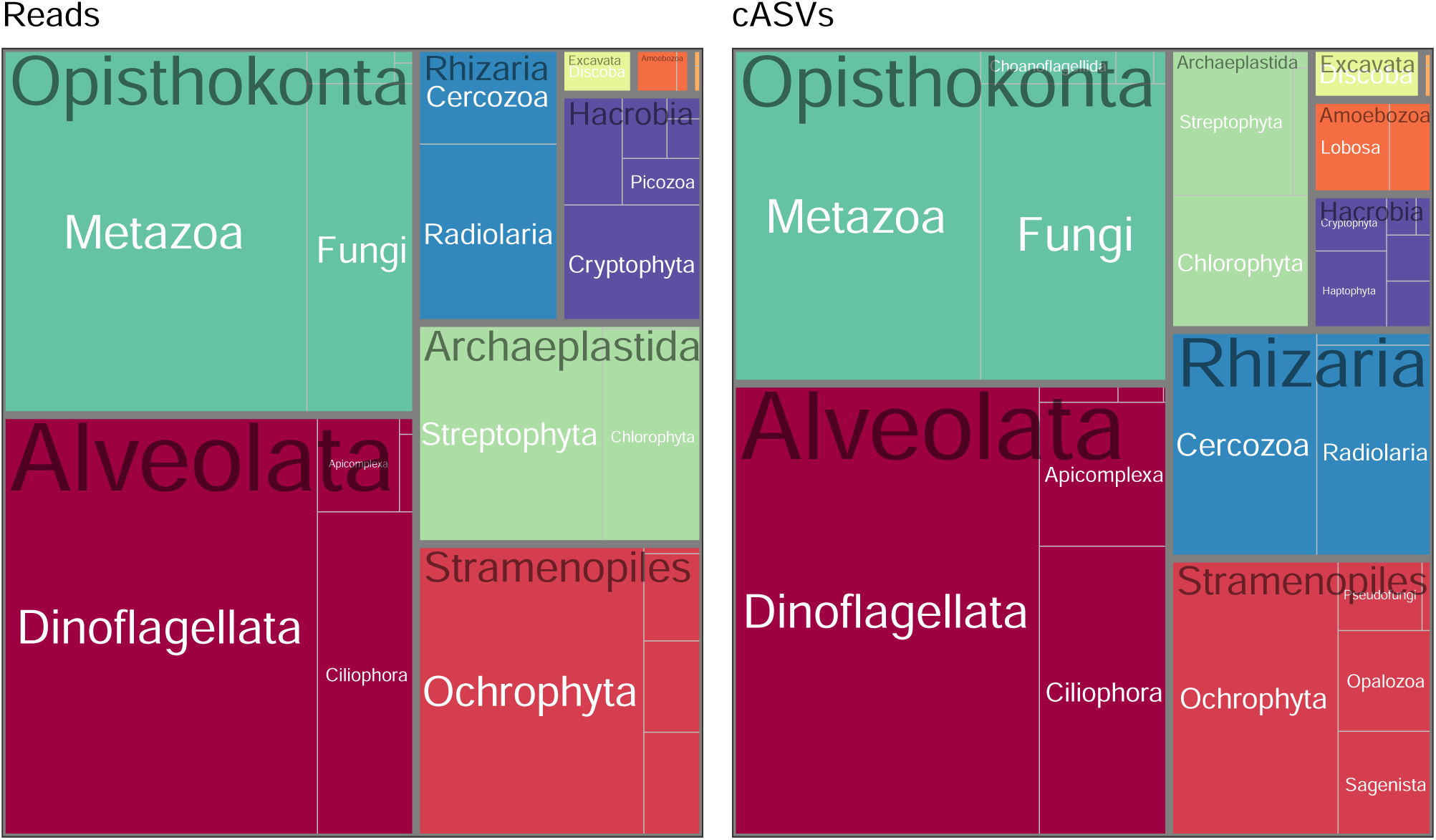
Treemaps of most abundant taxa (supergroup and division) for all datasets (V4 and V9) based on number of reads after normalization (left) or number of cASVs (right).

**Figure S5:**
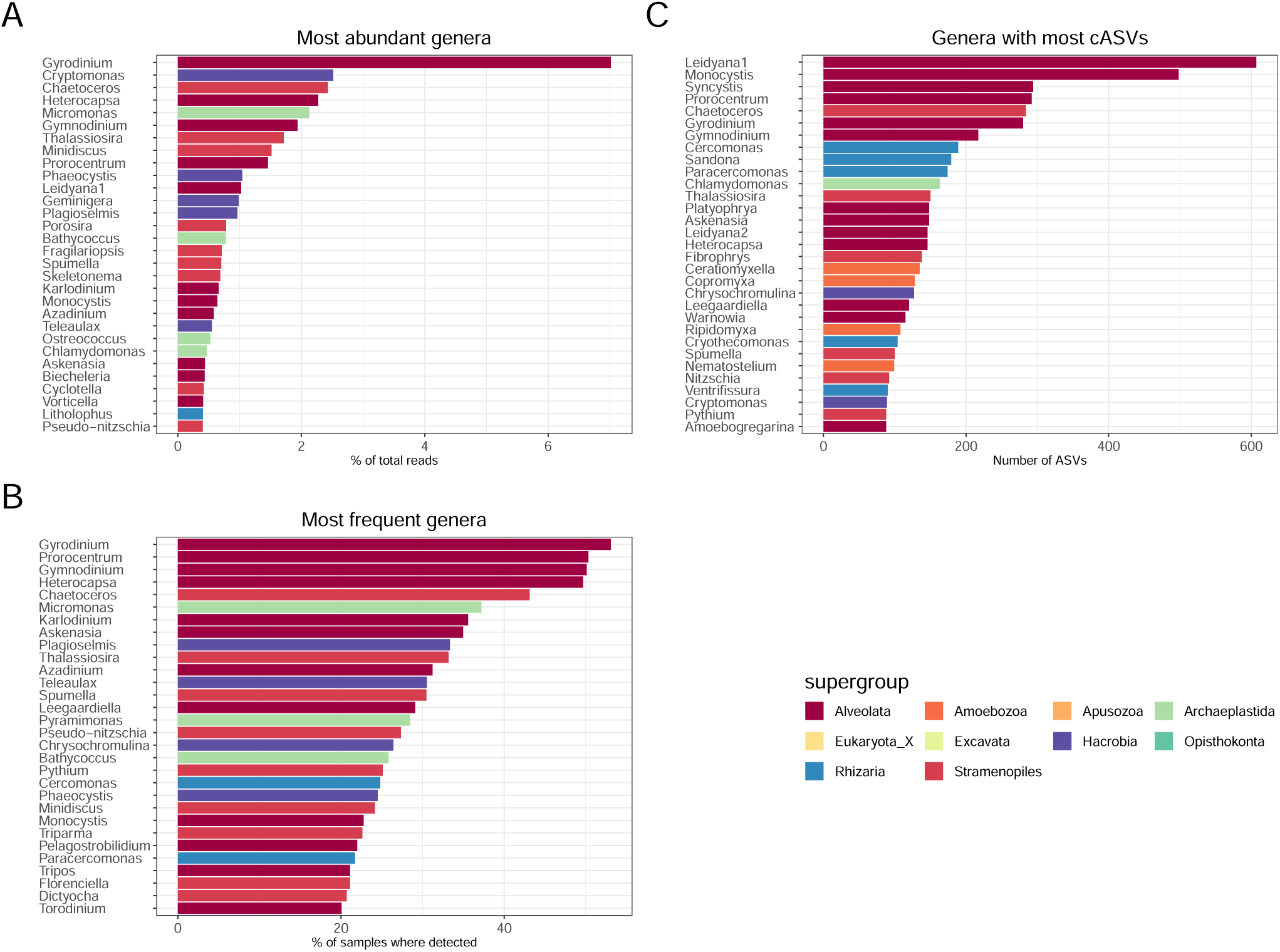
Protist genus analysis for for the V4 dataset after normalization. A. Most abundant genera. B. Most frequent genera. C. Genera with most cASVs.

**Figure S6:**
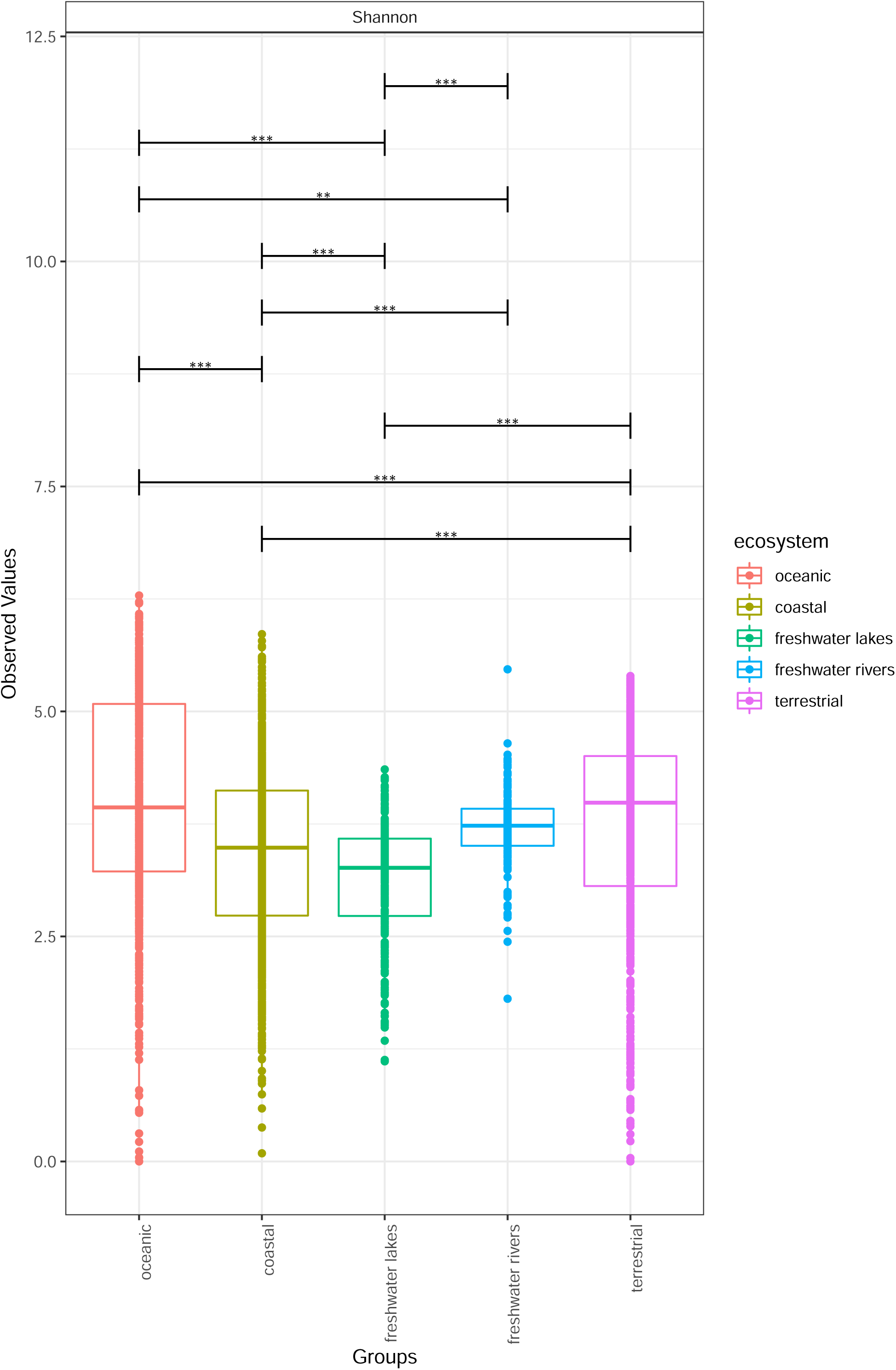
Protist V4 ASVs. Shannon’s diversity index as a function of the environment with significance (** p-value < 0.01, *** p-value < 0.001).

**Figure S7:**
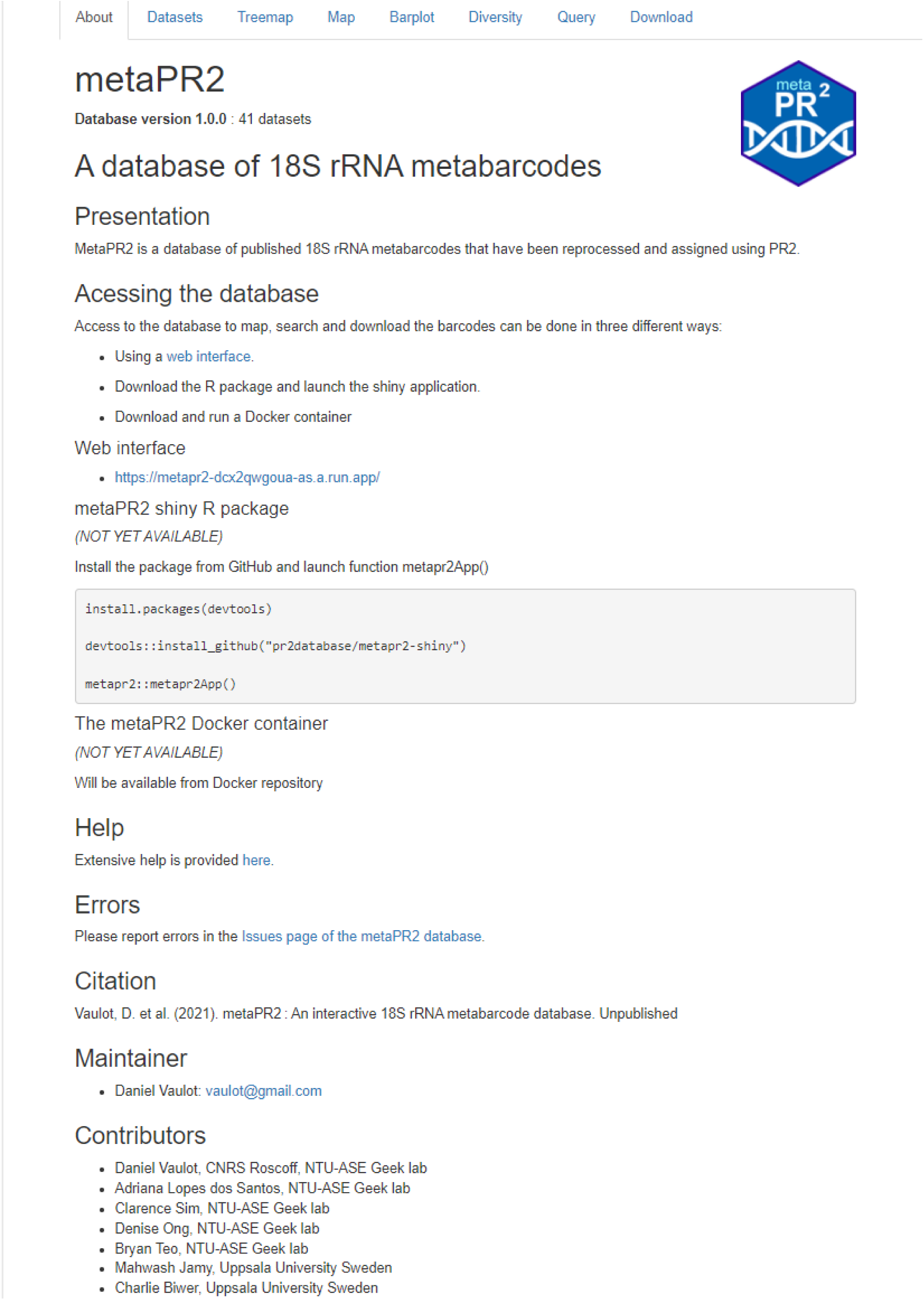
Shiny panel “about”.

**Figure S8:**
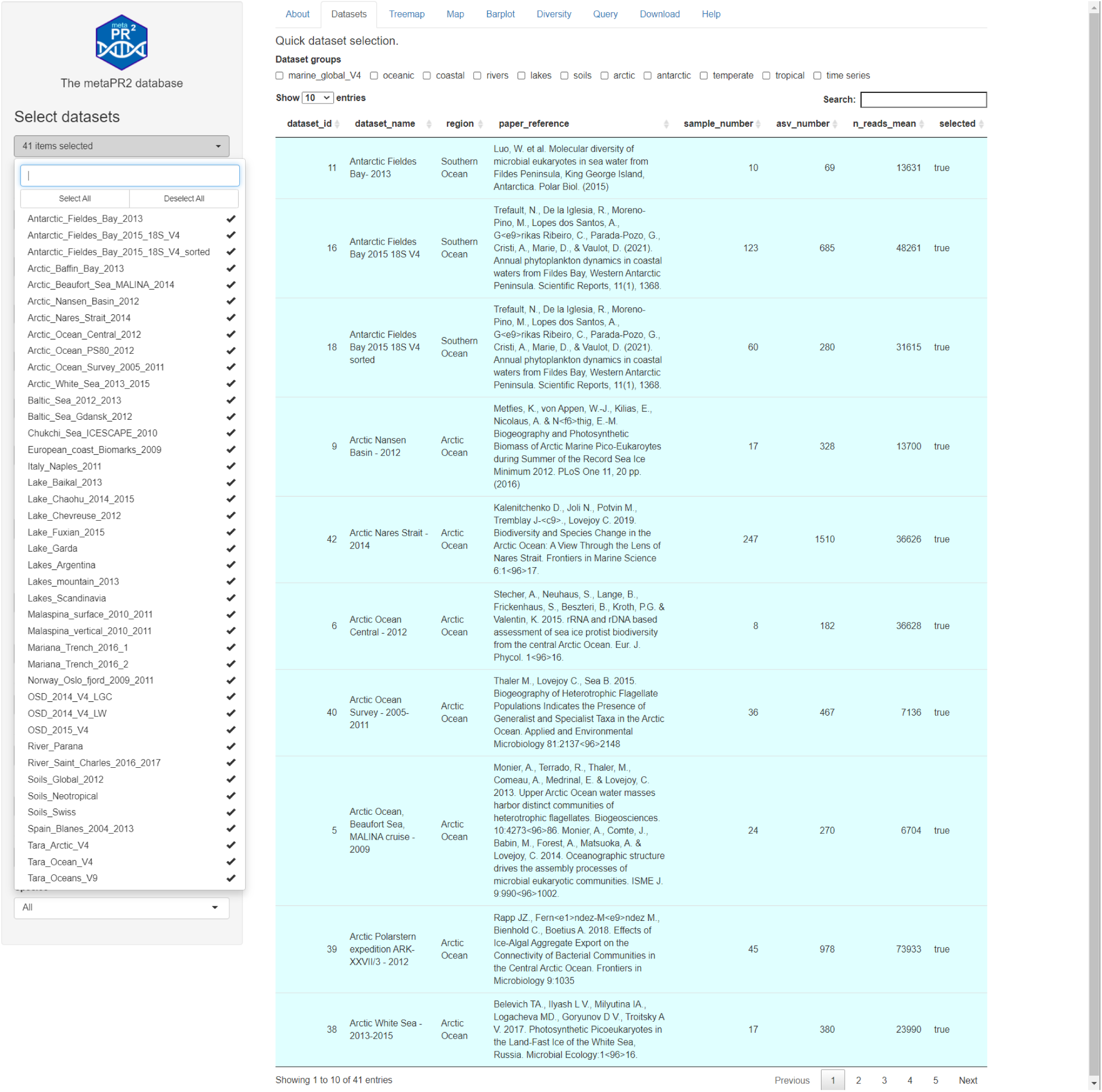
Shiny panel “datasets”.

**Figure S9:**
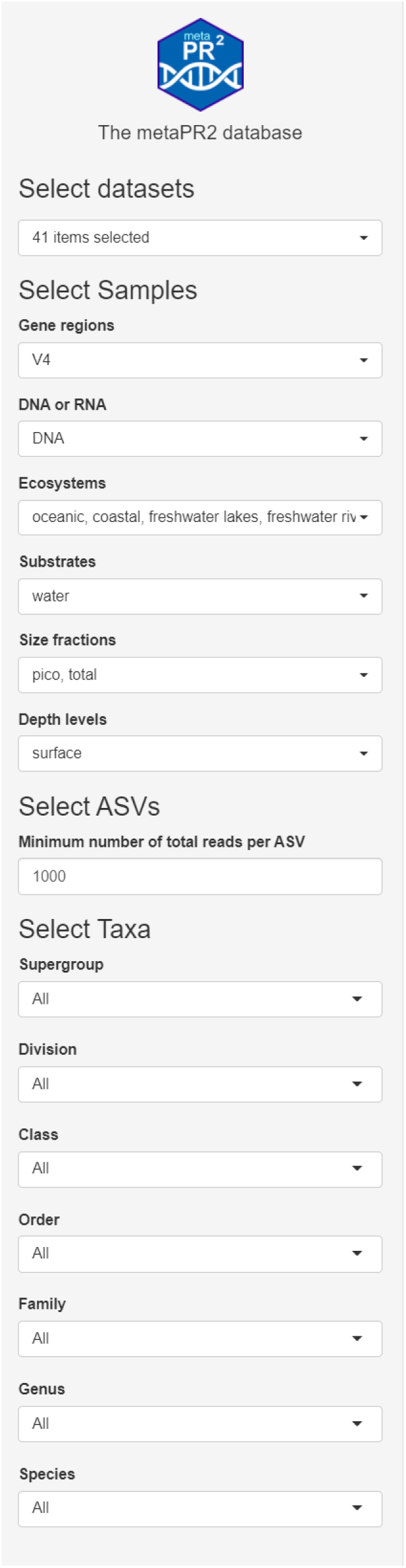
Shiny sample selection sidebar.

**Figure S10:**
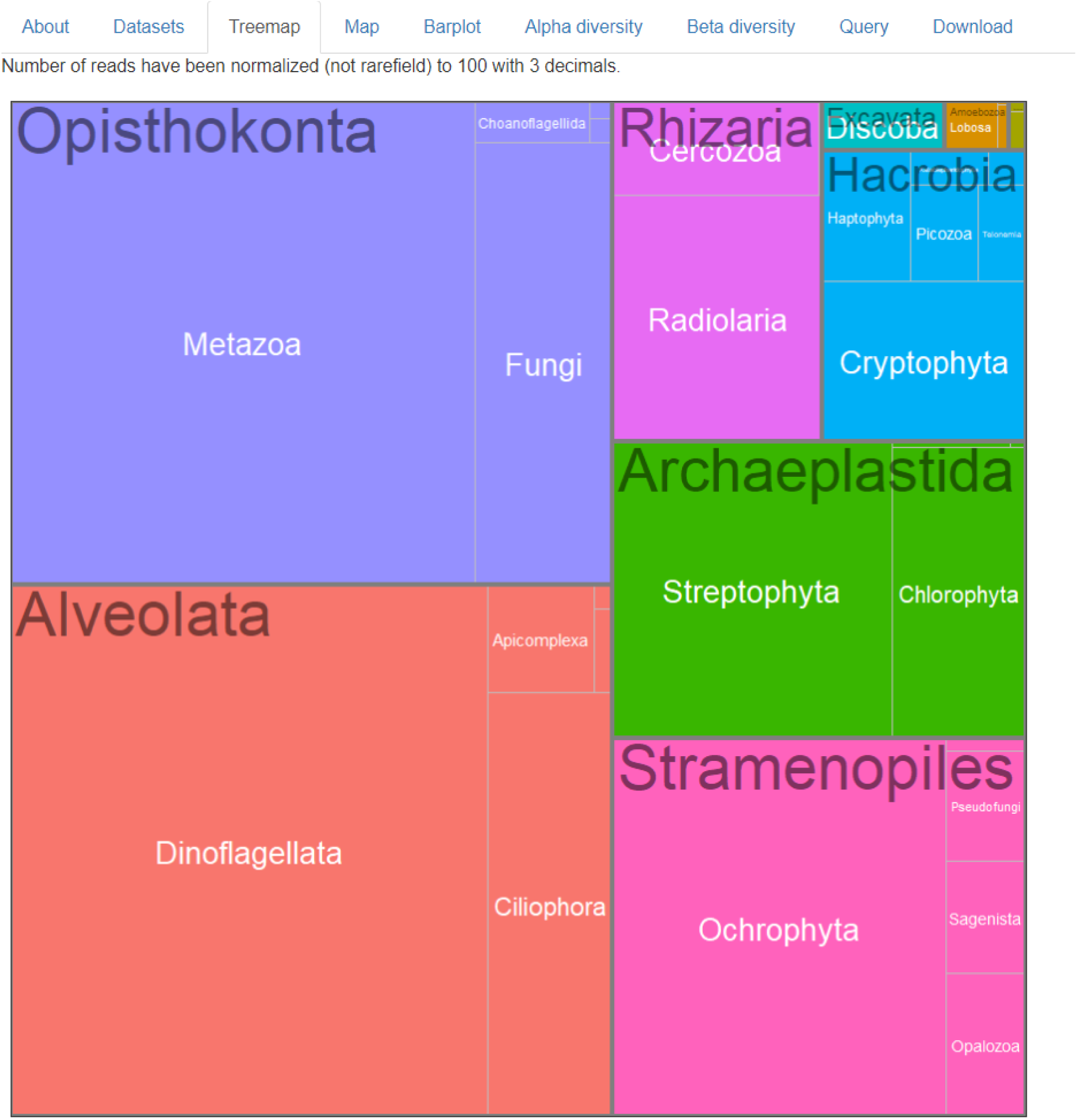
Shiny panel “treemap”.

**Figure S11:**
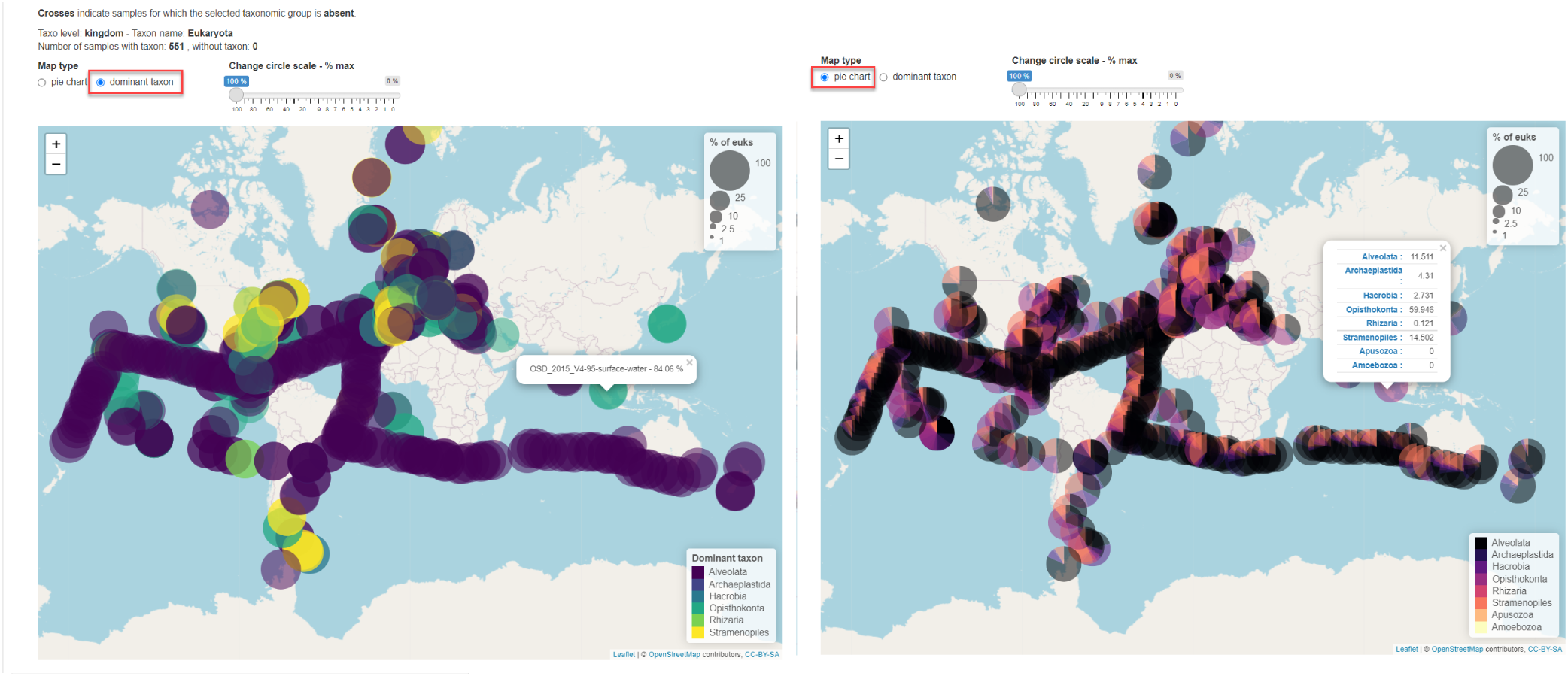
Shiny panel “map”.

**Figure S12:**
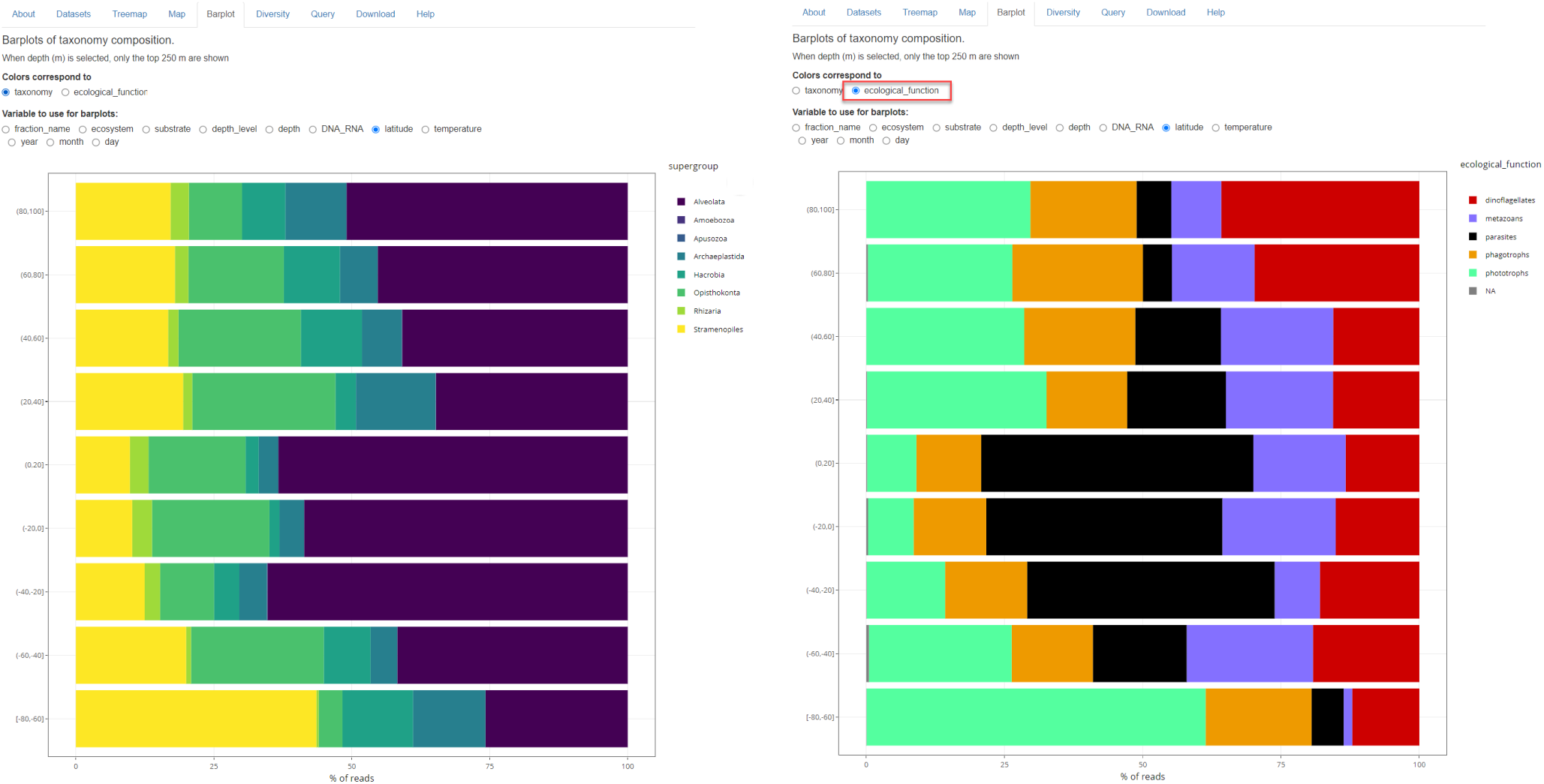
Shiny panel “barplot”.

**Figure S13:**
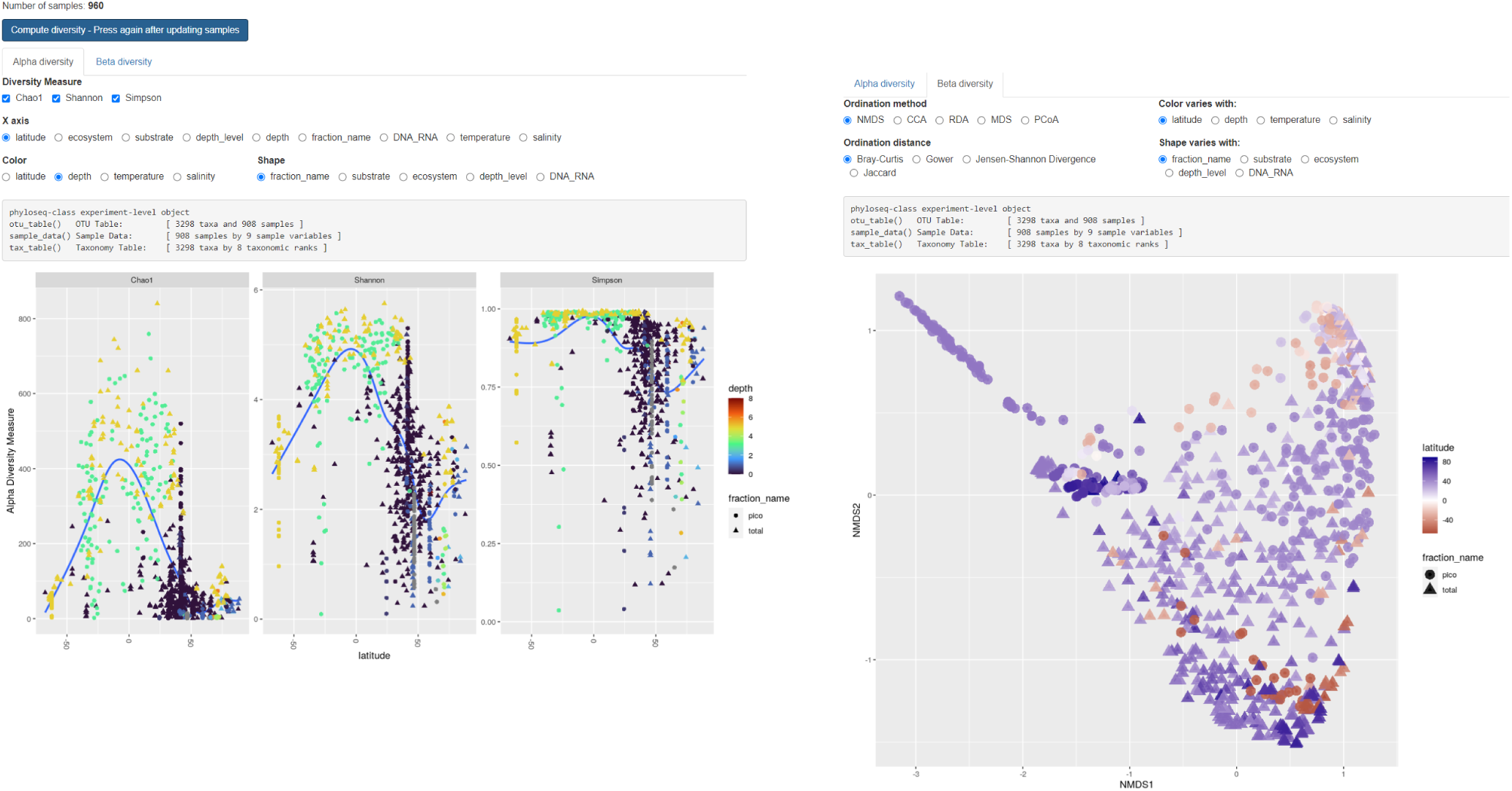
Shiny panel “diversity”.

**Figure S14:**
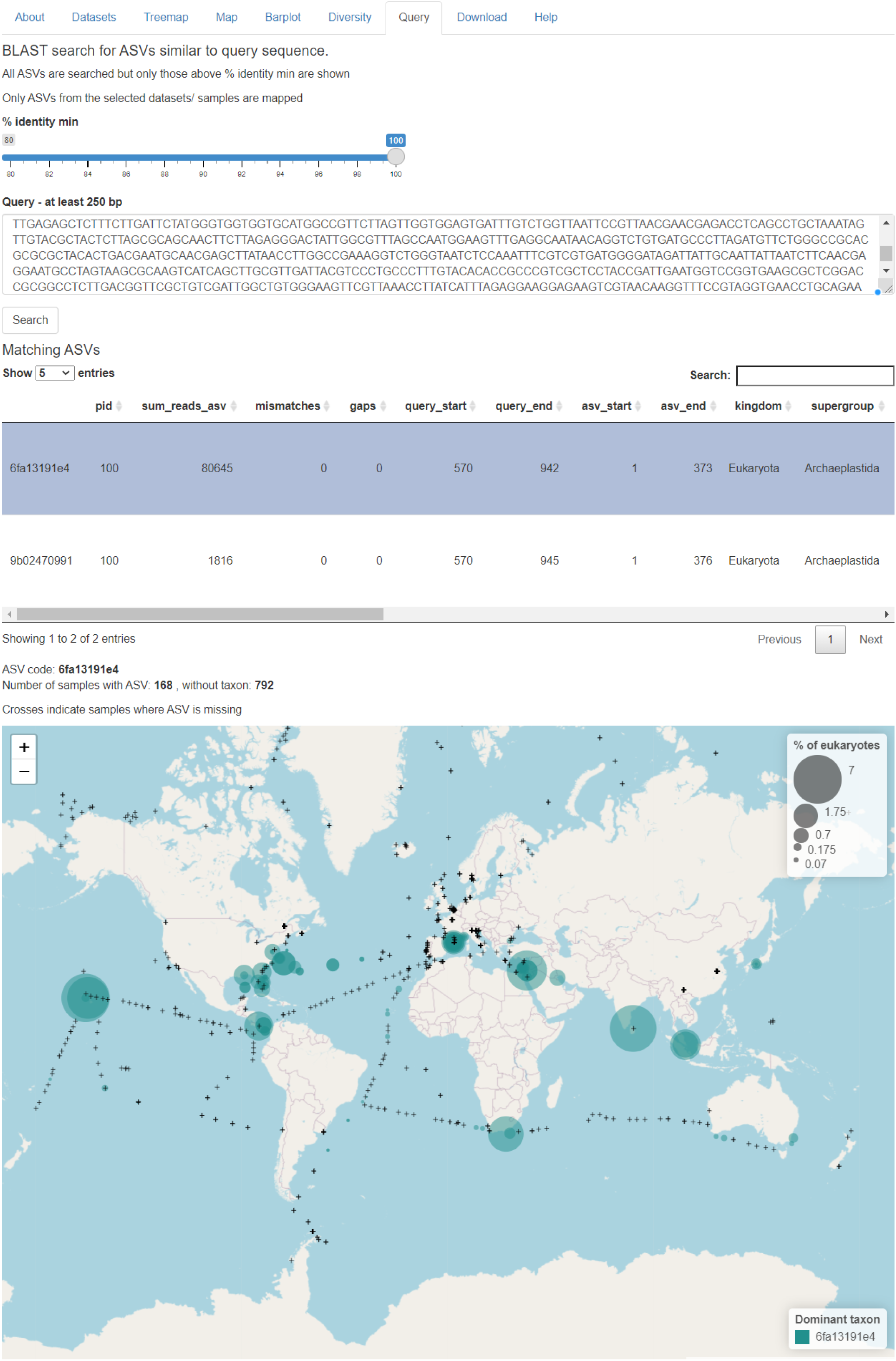
Shiny panel “query”.

**Figure S15:**
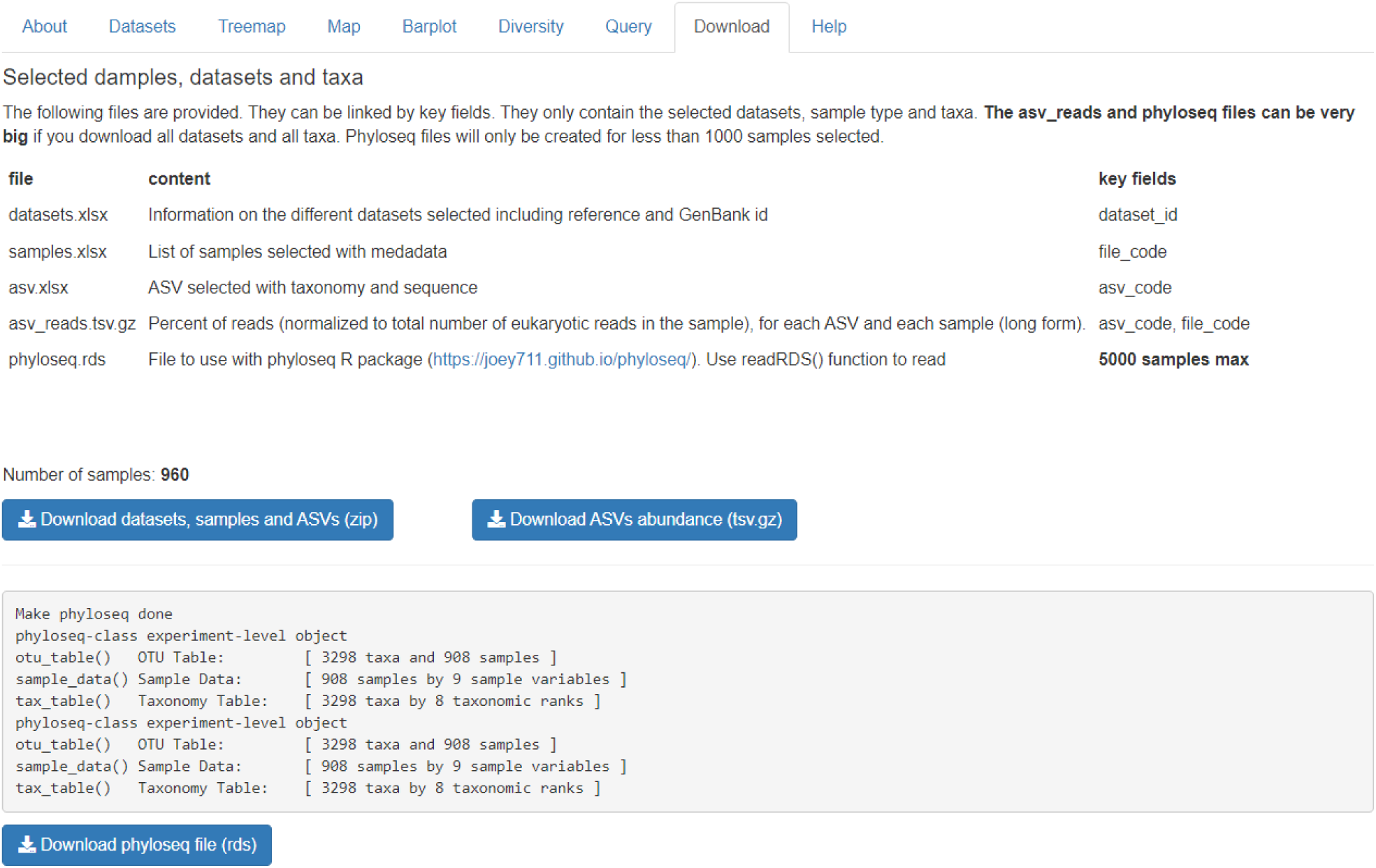
Shiny panel “download”.

